# Machine Learning prediction of cardiac resynchronisation therapy response from combination of clinical and model-driven data

**DOI:** 10.1101/2021.09.03.458464

**Authors:** Svyatoslav Khamzin, Arsenii Dokuchaev, Anastasia Bazhutina, Tatiana Chumarnaya, Stepan Zubarev, Tamara Lyubimtseva, Viktoria Lebedeva, Dmitry Lebedev, Viatcheslav Gurev, Olga Solovyova

## Abstract

**Background:** Up to 30%-50% of chronic heart failure patients who underwent cardiac resynchronization therapy (CRT) do not respond to the treatment. Therefore, patient stratification for CRT and optimization of CRT device settings remain a challenge.

**Objective:** The main goal of our study is to develop a predictive model of CRT outcome using a combination of clinical data recorded in patients before CRT and simulations of the response to biventricular (BiV) pacing in personalized computational models of the cardiac electrophysiology.

**Materials and Methods:** Retrospective data from 57 patients who underwent CRT device implantation was utilized. Positive response to CRT was defined by a 10% increase in the left ventricular ejection fraction in a year after implantation. For each patient, an anatomical model of the heart and torso was reconstructed from MRI and CT images and tailored to ECG recorded in the participant. The models were used to compute ventricular activation time, ECG duration and electrical dyssynchrony indices during intrinsic rhythm and BiV pacing from active poles of leads. For building a predictive model of CRT response, we used clinical data recorded before CRT device implantation together with model-derived biomarkers of ventricular excitation in the left bundle branch block mode of activation and under BiV stimulation. Several Machine Learning (ML) classifiers and feature selection algorithms were tested on the hybrid dataset, and the quality of predictors was assessed using the area under receiver operating curve (ROC AUC). The classifiers on the hybrid data were compared with ML models built on clinical data only.

**Results:** The best ML classifier utilizing a hybrid set of clinical and model-driven data demonstrated ROC AUC of 0.82, an accuracy of 0.82, sensitivity of 0.85, and specificity of 0.78, improving quality over that of ML predictors built on clinical data from much larger datasets. Distance from the LV pacing site to the post-infarction zone and ventricular activation characteristics under BiV pacing were shown as the most relevant model-driven features for CRT response classification.

**Conclusion:** Our results suggest that combination of clinical and model-driven data increases the accuracy of classification models for CRT outcomes.

## 1 Introduction

Cardiac resynchronization therapy (CRT) is one of the most effective non-pharmacological therapies for patients with chronic heart failure (CHF). It enhances the pumping function increasing the left-ventricular (LV) ejection fraction (EF), promotes reversed cardiac remodeling, and improves patients’ quality of life [1, 2]. Nevertheless, 30-50% of candidates for CRT have no significant improvement after implantation [3], which points to the importance of clarifying the criteria for patient selection and optimizing the implantation procedure itself.

Lack of response to CRT is a multifactorial problem associated with variability in individual characteristics, disease patterns, and treatment [4]. Combined assessment of multiple factors and individual patient characteristics can improve prediction of response to CRT. With increasing availability of electronic databases, Machine Learning (ML) provides an opportunity to perform such assessment, improving patient selection for therapy [5, 6]. Recent studies using ML techniques have achieved impressive results in preoperative clinical data analysis for selecting patients for CRT. Predictive models have been developed to estimate mortality or hospitalization risks from the baseline clinical parameters [7, 8, 9], to assess improvements in EF based on baseline indices and analysis of medical records [10] and to stratify patients by an unsupervised learning approach implementing ECG traces [11] and electrocardiography [12]. In a recent study [13], Feeny and co-authors showed 9 features (QRS morphology, QRS duration, New York Heart Association CHF classification, LV EF and end-diastolic diameter (EDD), sex, ischemic cardiomyopathy, atrial fibrillation, and epicardial LV lead) were sufficient to predict patient improvement with fairly high accuracy.

In addition to advances in ML approaches, significant progress has been made in computer modelling of the heart [14, 15]. Recent work has shown that patient-specific computer models based on 12-channel ECG and cardiac anatomy measurements are able to reproduce ventricular activation [16, 17, 18, 19]. Moreover, such models may be used to simulate the effect of CRT and study dyssynchrony characteristics [18, 20].

The focus of the present article is to predict the response to CRT using ML classification on combined clinical data with *in silico* simulations of biventricular (BiV) pacing in patient-specific models of ventricular electrical activity. We hypothesize that combination of simulation and clinical datasets could gain the predictive power of ML models of CRT response.

The study focuses on the following research aims: to assess the contribution of simulated indices derived from personalized electrophysiological modelling of BiV pacing to the accuracy of ML predictive models; and to define important clinical and model-derived features in the hybrid dataset for CRT response prediction.

## 2 Methods

The schematic outline of the research pipeline, including patient cohort selection, clinical indices’ acquisition, electrophysiological modelling, feature selection, and machine learning model training, is illustrated in Figure 1.

**Figure 1:**
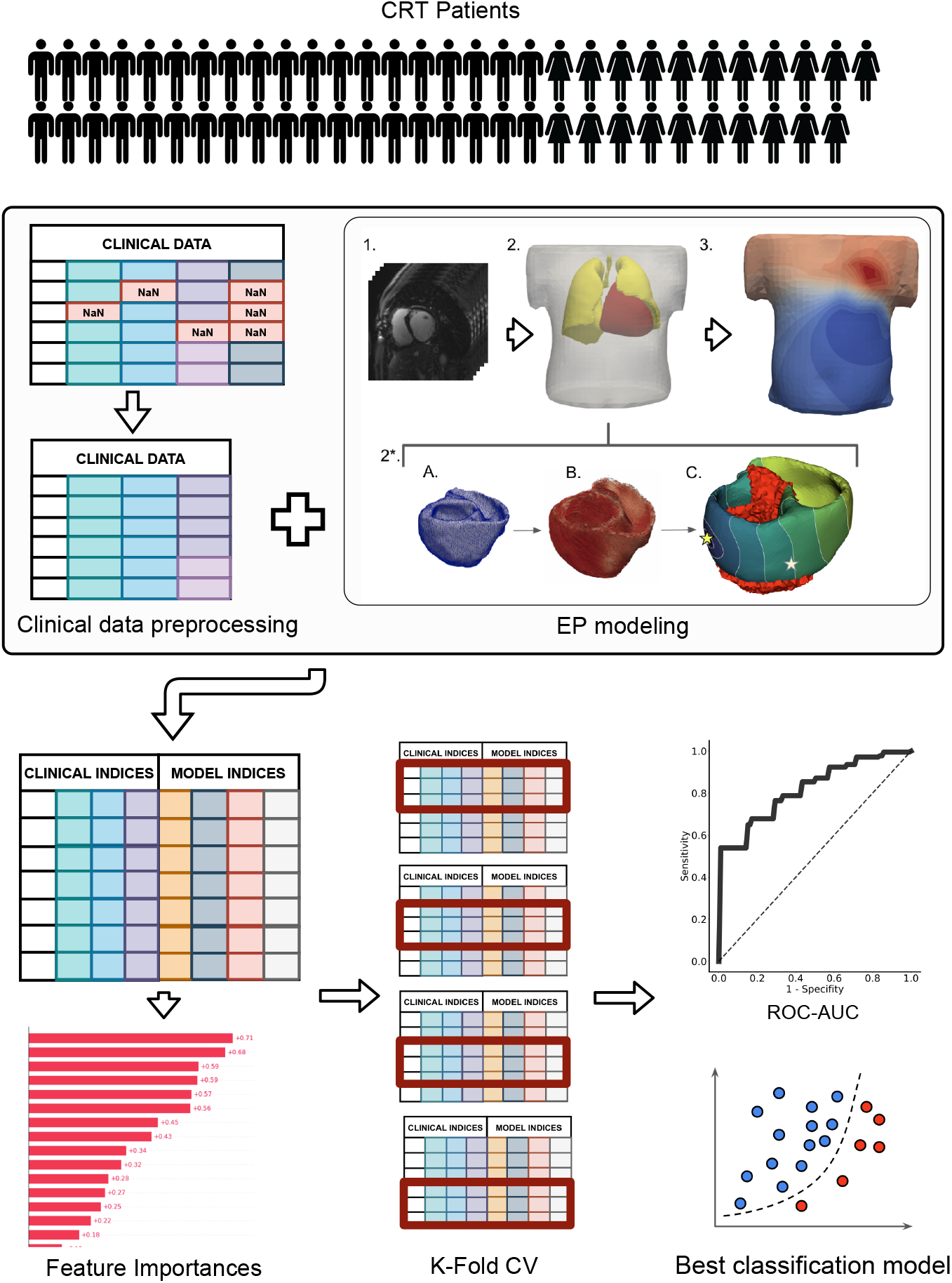
The schematic outline of the data analysis and machine learning pipeline. The pipeline included three major steps: I. CRT patient cohort assembling II. Preprocessing of clinical data and electrophysiological (EP) modelling III. Machine learning model development. In the clinical data preprocessing stage: features with missing values were excluded, non-categorical data were normalized by subtracting mean and dividing by standard deviation, collinear features were removed from the dataset by threshold > 0.85. EP modelling stage included: 1. CT data processing; 2. Segmentation of finite element meshes of the torso and lungs. 2*. Personalization of the heart model: a) Heart segmentation; b) Assignment of myocardial fibers [21]; c) Infarction scar/fibrosis assignment, pacing protocol selection (LBBB or BiV) and activation map calculation (stars indicate pacing sites), the infarction area is marked in red. 3. Calculation of torso potential map and ECG signals deriving.

### 2.1 Clinical data

#### 2.1.1 Study Population

In this retrospective non-randomized single-center observational study, we enrolled 57 CHF patients on optimal drug treatment who underwent CRT device implantation at Almazov National Medical Research Centre from August 2016 to August 2019. Participants signed approved inform consent. The study protocol was approved by the Institutional Ethical Committee.

The criteria for inclusion into the study were:

1. age over 18;
2. functional class (FC) II-IV of CHF according to the classification of the New York Heart Association (NYHA) at the outpatient stage of treatment;
3. LV EF ≤ 35% (Simpson);
4. QRS duration (QRSd) more than 120 ms;
5. sinus rhythm, left bundle branch block (LBBB);
6. optimal drug therapy.

The exclusion criteria were:

1. acute myocardial infarction, transient ischemic attack, acute cerebrovascular accident less than 3 months before the start of the study;
2. patients who were scheduled to undergo myocardial revascularization or heart transplantation during the observation period;
3. congenital and acquired defects, as well as heart tumors, LV aneurysm, if scheduled for surgery during the observation period;
4. active inflammatory and autoimmune diseases of the myocardium;
5. thyrotoxicosis at the time of inclusion in the study;
6. anemic syndrome: blood hemoglobin level less than 90 g/l;
7. diseases limiting life expectancy to less than 1 year.

#### 2.1.2 Data collection

Patients were evaluated before CRT device implantation and during the follow-up period of 12 months after implantation. Patients underwent investigation according to standard pro forma with some additional research methods appropriate for this study.

Standard research methods include:

- clinical examination (complaints, medical history, physical examination) - before CRT and 1 year after CRT;
- general blood test, biochemical blood test (glucose, potassium, sodium, creatinine, urea, total bilirubin and its fractions, total cholesterol, total protein, AST, ALT), general urinalysis - before CRT;
- 12-lead ECG - before and 1 year after CRT; ECG monitoring during CRT device programming and during the entire observation period;
- echocardiographic studies before and 1 year after CRT to assess LV reverse remodeling;
- stress tests to exclude/confirm coronary artery disease: stress echocardiography, bicycle ergometry or treadmill test, where clinically indicated;
- coronary angiography, where clinically indicated.

Additional research methods include:

- ECG recording in intrinsic rhythm and under BiV pacing, while programming the CRT device within 7 days after implantation.
- Electrocardiographic imaging using an Amycard system (Amycard, EP Solutions SA, Yverdon, Switzerland). Prior to ECG imaging, a maximum of 224 unipolar body surface mapping electrodes were placed on the patient’s torso, followed by computed tomography (CT) imaging of the heart and thorax (Somatom Definition 128, Siemens Healthcare, Germany). Subsequently, the electrodes were connected to the model 01C multichannel electrophysiology laboratory system (Amycard) for continuous ECG recordings during the pacing protocol. CT data were imported into Wave program version 2.14 (Amycard software) to reconstruct 3-dimensional geometry of the torso and heart. Finally, epi/endo ventricle models were manually built with marked active poles of RV and LV leads for bi-ventricular pacing simulations.
- MRI (MAGNETOM Trio A Tim 3 T, Siemens AG or INGENIA 1.5 T, Philips) with contrast (Gadovist or Magnevist) before CRT to detect structural lesions to the myocardium.
- Tissue Doppler echocardiography to record ventricle mechanical dyssynchrony. Analysis of interventricular dyssynchrony (IVD) and intraventricular dyssynchrony in the LV (LVD) was performed using biomarkers suggested by Yu and co-authors [22]. IVD was assessed by the time difference between the start of systolic flows into the aorta and the pulmonary trunk as measured by a pulse-wave Doppler, a value of less than 40 ms was taken as an IVD normal value. LVD was assessed using two biomarkers: dyssynchrony index defined as the temporal difference between the maximal and minimal peak systolic velocities between 12 LV segments (Tsmax–Ts min, 105 ms was taken as threshold normal value), and standard deviation in the peak systolic velocities for 12 LV segments (SD–12, 34 ms was taken as cutoff value). To determine the peak systolic velocities, the technique of colour tissue Doppler ultrasonography was used.

Baseline clinical data for the patients’ cohort is presented in Table 2.

**Table 1:** Model biomarkers

| Index | Definition | Explanation |
| --- | --- | --- |
| TAT, ms | $TAT = AT_{max} - AT_{min}$ | total ventricular activation time, $AT_{min}$ (ms) - activation start time, $AT_{max}$ (ms) - late activation time. |
| QRSd, ms | $QRSd = \max(T_S - T_Q)$ | maximum QRS complex duration in all leads - difference in the Q-S peaks time on the ECG signal. |
| $AT_{RV LV}$, ms | $AT_{RV LV} = AT_{max LV} - AT_{max RV}$ | interventricular dyssynchrony index - the difference between the time of late activation of LV and RV. |
| $mAT_{STLV}$ | $mAT_{STLV} = \frac{LVlat_{mean} - ST_{mean}}{TAT}$ | LV activation dyssynchrony index, where $LVlat_{mean}$ (ms)- average activation time of LV free wall, $ST_{mean}$ - average activation time of septum. |
| $IntAV_{STLV}$, ms | $IntAV_{STLV} = \int \left\vert \frac{AV_{LAT}(t)}{V_{LAT}} - \frac{AV_{ST}(t)}{V_{ST}} \right\vert dt$ | integral index of LV activation dyssynchrony, $AV_{ST}(t)$ - fraction of myocardial volume activated in septum at time t, $AV_{LAT}(t)$ - fraction of myocardial volume activated in free wall at time t, $V_{LAT}$ - free wall volume, $V_{ST}$ - septum volume. |

**Table 2:**
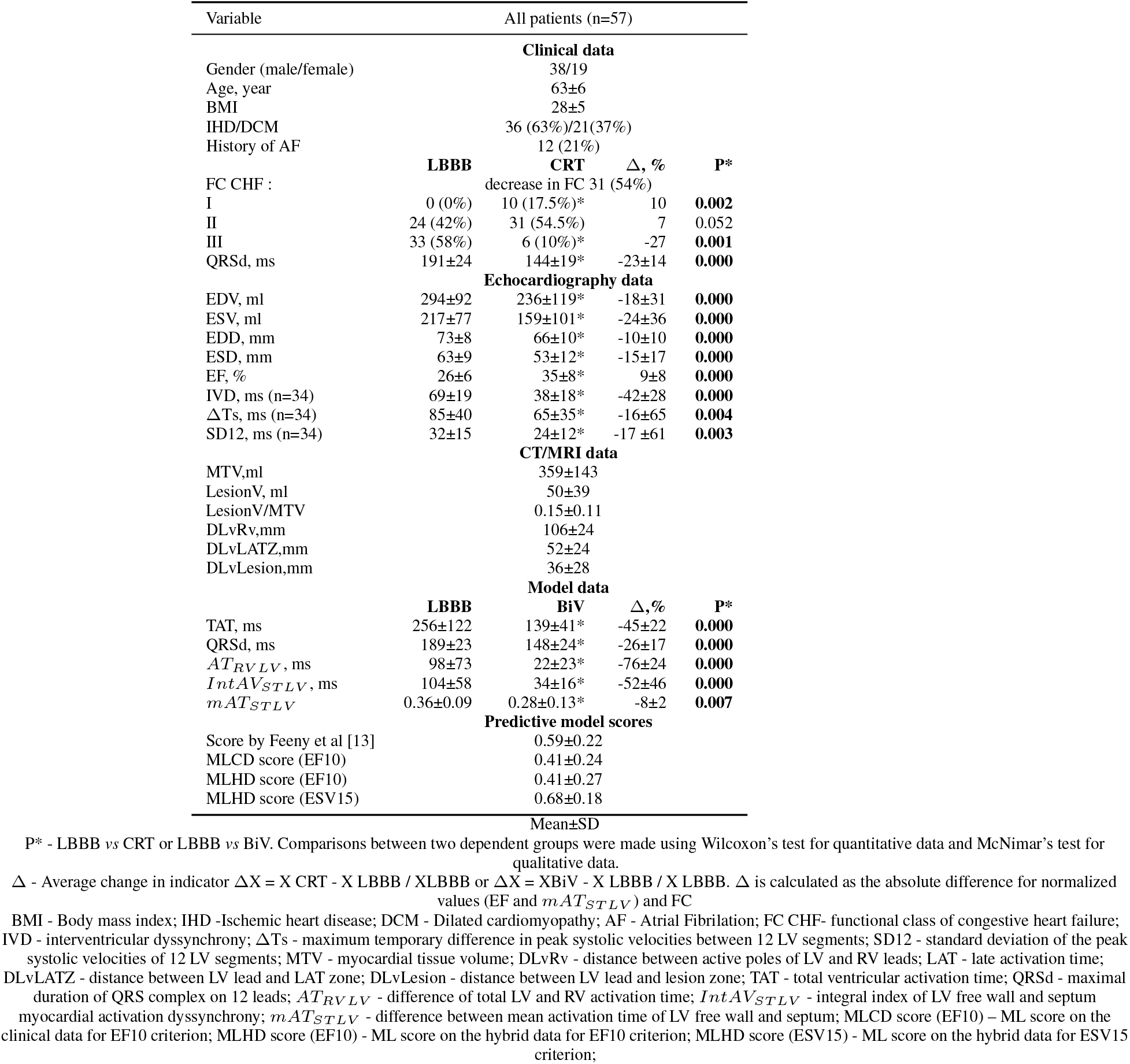
Clinical, imaging, model data and predictive model scores for the patient cohort.

### 2.2 Personalized ventricular models of electrophysiology

Patient-specific models were generated for each of the 57 cases. Semi-automatic CT segmentation approach helped to extract torso, lungs and ventricles. (Fig. 1 EP modelling, items 1-2). Finite-element meshes were smoothed, refined and merged. Average edge length was 15 mm for torso, 10 mm for lungs and 4 mm for heart.

Then, the LV myocardial tissue in the patient ventricular model was further annotated as either normal tissue or lesions according to the report of the expert who examined the patients’ MRI scans. The annotation was made using the conventional 17-segment American Heart Association (AHA) model of the LV [23], split into layers, endocardial, mid-myocardial and epicardial, in which damaged regions were highlighted. Two types of myocardial lesion were annotated based on the MRI report: ischemic and non-ischemic fibrosis. The regions of post-infarction scars were depicted as non-excitable, therefore such zones were excluded from model calculations. As for regions with myocardial fibrosis, the fibrotic tissue was assigned a low constant conductivity (1% of normal conductivity). Figure 2A shows a ventricular geometry model for patient #11 with a zone of intramural fibrosis (red) located in the septum (AHA segments 2,3,8,9).

**Figure 2:**
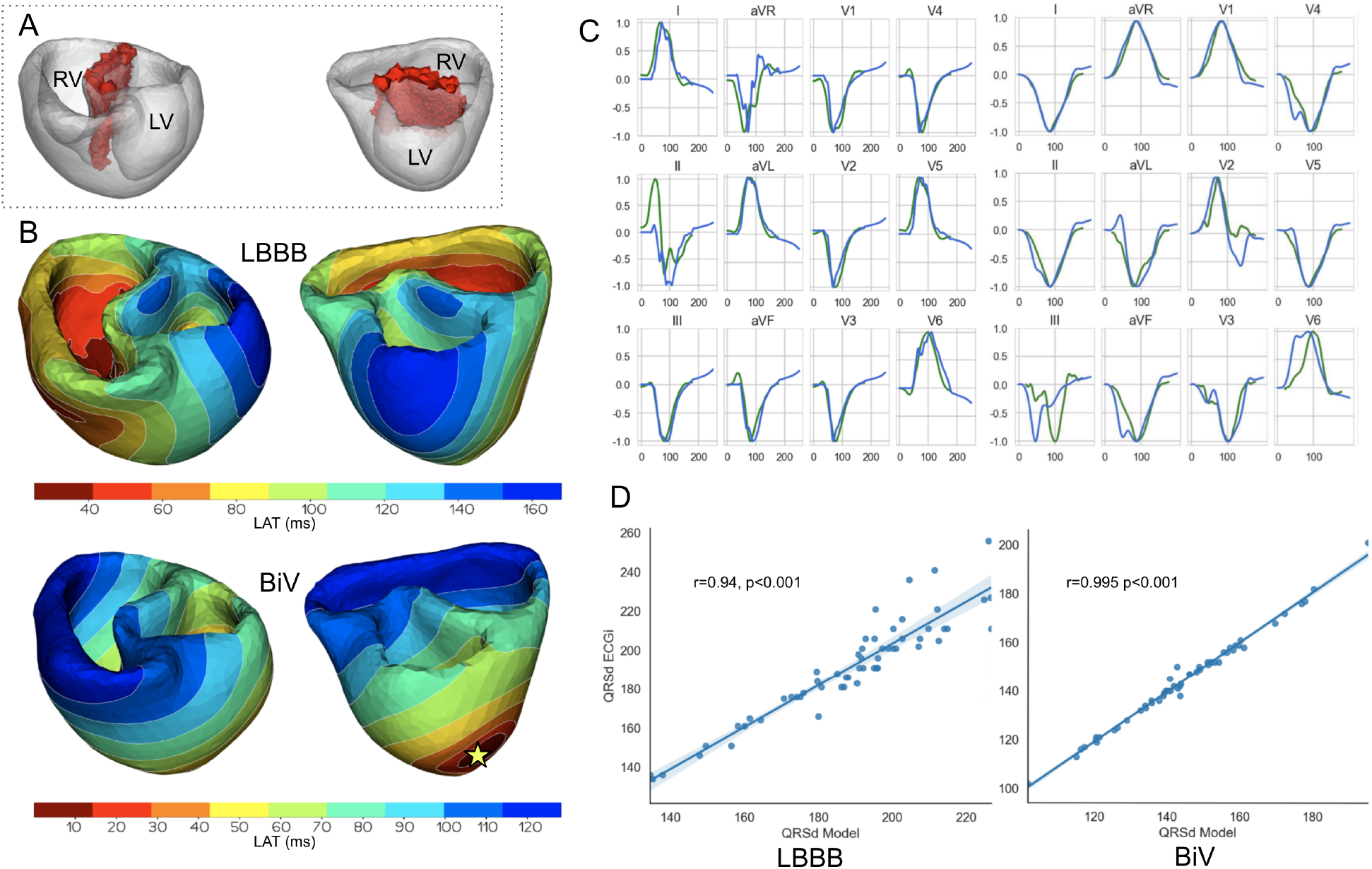
Model validation. Example of a personalized ventricular model for patient #11. A. Area of fibrosis in the interventricular septum (red zone). B. Comparison of model activation maps for LBBB (top) and BiV pacing (bottom). Star indicates LV pacing site for BiV pacing stimulation. The RV stimulating electrode was located in the apex of the surface. C. Calculated ECG signals (QRS complexes) for LBBB on the left and BiV pacing on the right. Green line - signals recorded in the clinic. Blue line - simulated signals. The amplitude of the QRS signals is normalized to the maximum values of the signals. D. Scatter plots showing correlations of QRS complex duration for 57 patients. Dots denote individual patients. The blue line is the regression line, Pearson correlation coefficient for LBBB is 0.94 (p<0.001) for BiV pacing is 0.99 (p<0.001).

For every patient-specific ventricular model, the electrical activity in the myocardium and ECG on the body-surface were simulated in several steps. We used an Eikonal model [24] to calculate the activation time at each point of the ventricular mesh. To assign fiber direction at every point of the myocardium, a rule-based approach was used [21]. Cardiac tissue was simulated as an anisotropic medium with a conductivity ratio of 4:1 along *vs* across the myocardial fibers providing the conduction velocity ratio 2:1 in the fiber and in the transverse direction, respectively. A global value of the conductivity along the fibers was set for the entire myocardial tissue and fitted against the clinical data as described in the next section.

Then, pre-computed cellular action potentials with time-shifts set according to the activation map were assigned to estimate time-dependent trans-membrane potential in the ventricles. For cellular action potential trace calculation, we used the TNNP06 model [25]. The resulting potential map was used to calculate the ECG by utilizing the pseudo-bidomain approach [26]. We defined the locations of 12-lead ECG electrodes on the torso surface and used simulated ECG signals for the analysis (Fig. 1 EP modelling, items 2-3).

#### 2.2.1 Simulation of LBBB activation pattern and BiV pacing

We calculated model-driven indices with reference to two types of ventricular pacing: LBBB activation pattern and BiV pacing.

For LBBB activation, RV sub-endocardial surface was annotated and a Purkinje network was generated using standard parameters from a Costabal model [27]. The right bundle branch was set at about 40 mm length, originating on the intraventricular septum, reaching the RV apex, and then splitting into the Purkinje fiber system of RV [28]. His system was isolated from the working myocardium and connected to it only at the ends of the Purkinje fibers through Purkinje-myocardial junction points (PMJs). We set the stimulation time in each PMJ according to the distance to the origin node divided by the conduction velocity in the His-Purkinje system, which we assumed to be 3 mm/ms [29].

The location of BiV pacing sites were derived from CT images. Active poles of RV and LV leads were annotated manually in Wave program version 2.14 (Amycard software). For BiV pacing simulations, we set a zero time delay between the RV and LV pacing sites as programmed in patients.

The activation time at the stimulation points for LBBB and BiV mode of activation was considered as a boundary condition for solving the Eikonal equation.

#### 2.2.2 Personalization of the electrophysiological models

Patient-specific ventricular models in both LBBB and BiV pacing protocols were fitted to reproduce individual data from recorded ECG with the ventricular pacing mode switched off (intrinsic rhythm with LBBB) and switched on (BiV pacing).

For each patient-specific model, we assumed a uniform conductivity in the myocardial tissue in the entire ventricles and solved an optimization problem to find a global conductivity parameter minimizing the discrepancy between simulated and clinical data recorded in the patient in either LBBB or BiV stimulation protocol independently. The post-infarction scar regions were excluded from the model tissue, and a low conductivity of 1% of normal value was assigned to the fibrotic tissue regions when solving the optimization problem. The global conductivity parameter was fitted to minimize the difference between the means of simulated and clinically measured QRSd from the 12-lead ECG recorded in the patient. We used L-BFGS-B algorithm built into SciPy.minimize routine to handle optimization in the model.

Figure 2 B, C shows the personalization results for patient #11 with intramural fibrosis located in the septum (AHA segments 2, 3, 8, 9 depicted in red in Figure 2 A). Although the model parameters were fitted to minimize the difference between the means of simulated and clinical QRSd, the morphology of the simulated QRS complexes corresponded well with clinical ones (see Figure 2C, blue lines show model signals, green lines show recorded clinical signals). The scatter plots in Figure 2D demonstrate high correlations between simulated and clinical QRSd for both LBBB and BiV modes. The higher correlation coefficient for BiV pacing is explained by precise positioning of the pacing sites derived from CT imaging data, while for LBBB we used a synthetic model of ventricular activation that does not reflect the morpho-anatomical characteristics of the RV conduction system in a particular patient.

#### 2.2.3 Model-derived biomarkers of myocardial lesions, pacing site location, and myocardial electrical activity

Our patient-specific models allowed us to identify several clinically important features affecting ventricular activation. The first group of model-derived indices are based on CT and MRI data coupled with electrophysiology model simulations. Using a digital ventricular model, we were able to define the volume of post-infarction scar and non-ischemic fibrosis and their size relative to the myocardial tissue volume. Knowing RV and LV active poles positions, we measured distance between them. Furthermore, distances from LV pacing site to the lesion zone and to the area of late activation time (LAT) under intrinsic rhythm were calculated. To measure the spacing within the computational mesh, we solved an isotropic Eikonal equation showing electrophysiological distance between the model points.

The second group of model-derived indices was calculated in LBBB and BiV mode of myocardial activation, these used as predictions of the effects of BiV pacing on electrical synchronization. Table 1 presents a complete list of the simulated characteristics with related definitions and formulas. We simulated the time activation map for both chambers and 12-lead ECG and calculated the following biomarkers derived from the time-dependent signals for further analysis of CRT response: total ventricular activation time (TAT), maximum QRS complex duration, difference between the total LV and RV activation times (*AT_LV RV_*), relative difference between the mean activation times of LV free wall and septum (*mAT_STLV_*), integral index of LV free wall and septum myocardial volume activation (*IntAV_STLV_*). The last three indices characterizing inter- and intraventricular electrical dyssynchrony of myocardial activation were used in work of Villongco and co-authors [20].

The average values of all model-derived indices for our enrolled cohort are shown in Table 2.

### 2.3 A predictive model of response to CRT based on preoperative clinical data and electrophysiology model simulations

For classifier development, we applied several supervised machine learning (ML) approaches to identify an optimal set of features and learning algorithm combination showing the best performance characteristics on hybrid data for our patient cohort. The hybrid dataset for building the classifier contained clinical and model-derived features as described above. At the preprocessing step, features with missing values were excluded. Non-categorical data were normalized by subtracting the mean and dividing by standard deviation. Collinear features were also removed from the dataset by threshold > 0.85.

Several criteria for CRT response definition were used for classification. The primary criterion for responders was more than 10% increase in LV EF (EF10) [13]. The following criteria were also considered: a reduction in ESV>15% (ESV15, see [30, 31]); a 5 and 15% increase in LV EF [13], and combined EF10 and ESV15.

We evaluated several classification algorithms: logistic regression (LR), linear discriminant analysis (LDA), support vector machine (SVM) with linear kernel, random forest (RF) classifier; each evaluated in combination with three different feature sets obtained by feature selection methods. The following algorithms were used for feature selection: random forest mean decrease accuracy (MDA), univariate statistical testing (UST, two-sample t-test for continuous variables and chi-squared test for categorical variables), and L1-based feature selection (L1, based on weights of LR). Features were selected in a cross-validation loop for each subset. The top 8 features chosen by the algorithms were used to construct the classifiers.

Feature selection and training of classification algorithms was done using a Leave-One-Out cross-validation loop. Within the loop, the ML classifier score for each test fold (hear each consisting of just one observation) was calculated. These ML scores were combined into one set to build the receiver operating characteristic (ROC) curve and to calculate the area under the ROC curve (AUC). The highest-performing combination of the classifier with feature selecting algorithm was chosen to develop the final classifier.

In addition, we used repeated stratified five-fold cross-validation with 1000 iteration in order to be confident in assessing the quality of the classifier. We quantified the classification performance of each feature set–algorithm combination with ROC AUC across all folds and iterations. The classifier with the highest ROC AUC was selected as the final classifier for our hybrid dataset.

#### 2.3.1 Software

Cardiac electrophysiology was modeled with the help of software written at the Institute of Immunology and Physiology UB RAS based on FENICS library (for solving PDE problems) [32] and VTK (for working with meshes). For the machine learning pipeline (see Fig. 1): classifier development, statistical modelling, feature selection, cross validation, and ROC-AUC calculation we used the sklearn library.

#### 2.3.2 Statistics

Detailed analysis was performed using the IBM SPSS Statistics 23.0.0.0 software package (USA). For qualitative data, the frequency and percentage of total patients in the cohort were calculated. Quantitative data are presented as mean ± standard deviation. Comparisons between two dependent groups were made using Wilcoxon’s test for quantitative data and McNimar’s test for qualitative data. Nonparametric Friedman’s two-way ANOVA was applied to compare related groups. Comparison between two independent groups was carried out using the Mann-Whitney test for quantitative data and Pearson’s chi-square test for qualitative data. Feature dependence was assessed using the Spearman rank correlation test. The critical level of statistical significance was taken equal to 0.05.

## 3 Results

Below, changes in feature values under BiV pacing against the LBBB baseline are expressed in relative units. For clinical characteristics, ΔX*_CRT_* =(X*_CRT_*-X*_LBBB_*)/X*_LBBB_*, where X*_LBBB_* and X*_CRT_* are feature values before and after CRT device implantation, respectively. For simulated indices, ΔX*_BiV_* =(X*_BiV_* -X*_LBBB_*)/X*_LBBB_*, where X*_LBBB_* and X*_BiV_* are feature values in the LBBB activation mode and BiV pacing. For the indices that are initially showed in relative units, e.g. EF and mAT*_STLV_*, the response to pacing is expressed as an absolute increment in the LBBB value: ΔEF_*CRT*_ =EF_*CRT*_-EF_*LBBB*_, and ΔmAT_*STLV BiV*_ =mAT_*STLV BiV*_ -mAT_*STLV LBBB*_.

For some features, we also used normalized characteristics, i.e. the ratio of the value to the myocardial tissue volume (MTV). Note, MTV should not be confused with the volume of the cavity inside the ventricle. We can evaluate MTV using a digital model of the ventricular geometry based on CT images. When using such normalization, the volume of the preserved myocardium only is taken into account, without allowing for the myocardial lesions. Since the scar tissue is not excited and does not contract, it is excluded from the volume of the active ventricular myocardium. Normalized features give a characteristic’s values per unit volume of the myocardium (analogous to the values per mass unit of the myocardium). For instance, TAT/MTV indirectly reflects a reciprocal value of the average velocity of myocardium activation in the ventricles.

### 3.1 Analysis of clinical data before and after CRT device implantation and model simulations in LBBB and BiV pacing. Responders versus nonresponders

A summary of clinical data statistics before and after CRT device implantation, CT/MRI derived data and model-driven biomarkers in LBBB and BiV pacing in the entire patient cohort is presented in Table 2. It is clearly demonstrated that all clinical indicators of the CRT outcome show an average positive response in the entire cohort. On average, QRSd is decreased by 23±14%, EDV and ESV are decreased by -18±31% and 24±36% respectively; EF is increased by 9±8%; NYHA functional class is decreased at least 1 point in 54% of patients. Mechanical dyssynchrony indices characterising asynchrony in inter- and intra-ventricular contraction show prominent average decrease in the patient cohort.

In consistency with the clinical data, the model simulations also demonstrate an average positive outcome of BiV pacing as compared to the LBBB activation (Fig. 3). Average TAT and QRSd are decresed by 45±22% and 26±17% respectively, and the latter is in good agreement with the clinical effect of BiV pacing on the QRSd. The electrical dyssynchrony indices also reveal a prominent decrease in the population of models, with the highest reduction in the inter-ventricular dyssynchrony index AT*_RV LV_* by 76±24%.

**Figure 3:**
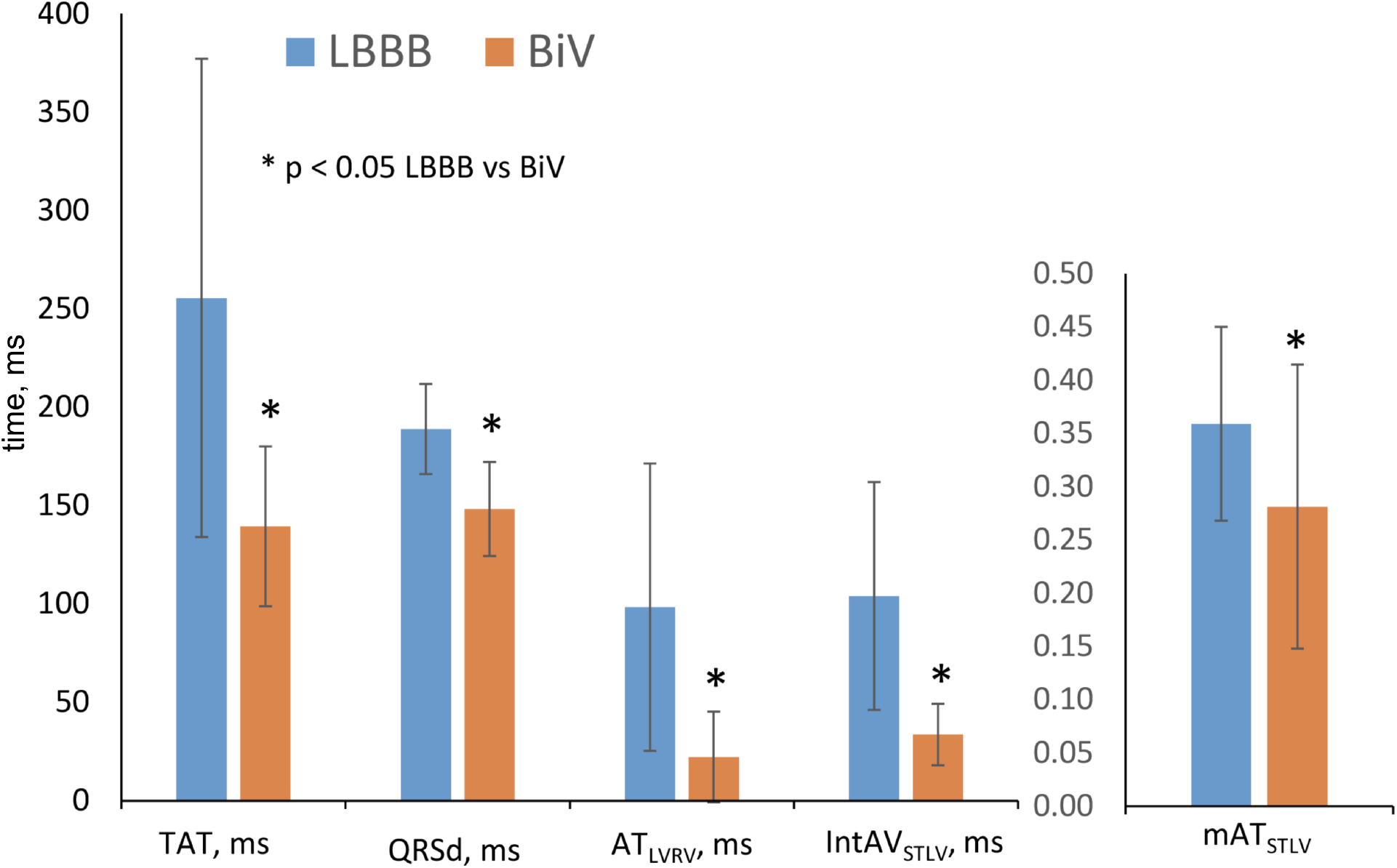
Simulation features in the LBBB activation mode and under BiV pacing. Bar indicates mean. Error bar is SD. Comparisons between two dependent groups (LBBB vs BiV) were made using Wilcoxon’s test.

However, we can see a high variability in the biomarker responses to the BIV pacing in both the clinical and simulated data. Coefficient of variation (SD/mean) in the relative change of some features is higher than 100% (e.g. see ΔEDV, ΔESV, and Δ for mechanical inter-ventricular dyssynchrony indices in Table 2), suggesting a significantly nonuniform output among the patients. Here, we have used several conventional criteria to classify responders and nonresponders to CRT in the patient cohort based on clinical data on the post-operative LV reversed remodeling. Primary classification was defined by a higher than 10% increase in the LV EF for responders (referred hereafter as EF10 criterion). This criterion was used in clinical studies, and allowed us to compare qualitatively the results of our predictive models for CRT response with the findings reported recently by Feeny and co-authors [13]. Surprisingly, the 10% cutoff for EF improvement in responders is close to the average EF increase observed in our patient cohort. Classification results based on other CRT response definitions are described in the Supplementary Materials and discussed in the Discussion Section.

Table 3 summarizes the clinical and model-derived variables in the groups with or without LV EF improvement according to the EF10 criterion. In our patient cohort, 23 (40%) patients demonstrated an improved EF (referred to as CRT responders) and 34 (60%) patients were classified as nonresponders. The ratio seems biased towards nonrespondents, but we have intentionally raised the LV EF improvement threshold in order to be more confident in predicting true positive responses. Average EF is raised in both groups, and the increase is significantly higher in the responders versus nonresponders (17±5% vs 3±5%, respectively). The EF improvement after CRT is accompanied by a prominent ESV reduction by 47±19% in the responder group against an insignificant diminishing by 9±37% in nonresponders. Similarly, a much higher average EDV reduction is seen in the responders due to LV postoperative reverse remodeling after CRT. Although the average QRSd is decreased, no statistical significance between the groups was found. No difference in the CRT effect on the mechanical dyssynchrony indices was found as well.

**Table 3:**
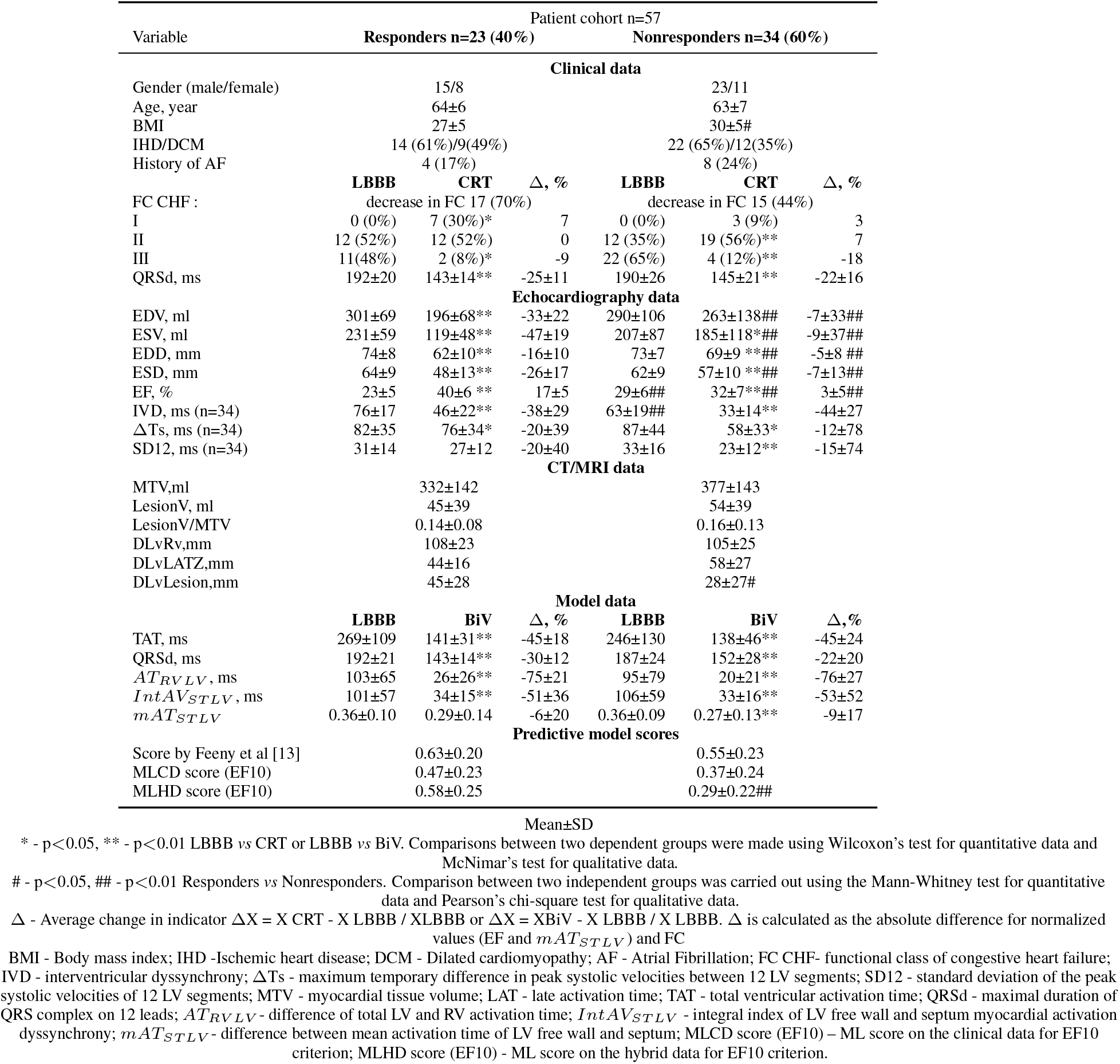
Clinical, imaging, model data and predictive model scores for responders and nonresponders defined by EF10 criterion.

In consistency with the clinical data, the model simulations revealed a decrease in both TAT and QRSd under BiV mode in each group. Meanwhile, no difference in the relative TAT and QRSd decrease between groups was observed (Table 3). Similarly, no significant difference was found between the groups in the effect of BIV pacing on the electrical dyssychrony indices calculated in the models. Using the LV geometry model built on the CT and MRI, we were able to compare the dimensions of lesions in LV myocardial tissue, and interlead spacing between active poles and distance from the LV electrode pacing site to the lesion zone as well (Table 3). We figured out no difference in the relative volume of the lesions in LV myocardial tissue between the groups. At the same time, we found a shorter distance from the LV pacing site to the lesion zone in the nonresponder group, suggesting less effective pacing of the normal tissue. Interlead spacing does not statistically differ between groups.

It is of note that most of the individual biomarkers in the intrinsic LBBB activation pattern derived from either clinical, or CT/MRI, or simulated data do not show a significant difference in the distribution between the responder and nonresponder groups. This means that no one single index could be considered as a diagnostic feature for preoperative classification (Table 3). Among the pre-operative clinical data, two features, i.e. LV EF*_LBBB_* and the inter-ventricular mechanical dyssynchrony index IVD*_LBBB_*, displayed differences between the groups classified according to the EF10 criterion. Here, LV EF demonstrated a bit higher average value along with a bit lower value of IVD in the nonresponders than in responders. This is consistent with a low negative correlation between EF*_LBBB_* before and ΔEF*_CRT_* after implantation (r=-0.48, p=0.031, see Figure 9 in the Supplementary Materials). However, high EF variation in each group comparable with the difference between the group averages did not allow us to find a valid threshold separating the groups. The average accuracy of the Logistic Regression classification with One-Leave-Out cross-validation based on EF*_LBBB_* was only 0.62 with rather low values of both sensitivity at 0.69 and specificity at 0.55. A low positive correlation was also found between IDV*_LBBB_* and ΔEF*_CRT_* (r=0.32, p=0.029), suggesting its possible predictive power for CRT response. However, we did not have IDV and other mechanical dyssynchrony indices for all 57 patients in our cohort and, therefore, decided against using them in further analysis (see Limitations Sec.).

Although in this proof-of-the-concept study some data used for model building were recorded after operation, we consider all CT/MRI and model-derived features as potentially pre-operative because all of them can actually be assessed before operation (see also Limitations Sec. on this issue). Among the data derived from CT/MRI, we found only one feature showing a significant difference between the responders and nonresponders: the distance between the LV pacing site and the lesion zone (Table 3). However, this index did not show a significant correlation with either ΔEF*_CRT_* or ΔESV*_CRT_* (see Figure 9 in the Supplementary Materials) as well. In consistency with the absence of difference between clinical QRSd in responders and nonresponders, none of the simulated electrophysiological biomarkers showed any significant difference between groups both in the LBBB mode of activation and under BiV pacing either (Table 3), which also did not allow them to be considered as individual classifying features.

Our dataset analysis suggested a hypothesis that the only combination of the clinical and MRI/CT derived biomarkers that can be evaluated before operation together with predictions on the BiV response simulated using a personalized ventricular model may increase the predictive power of such a hybrid dataset for patient classification.

### 3.2 Correlations within the input (pre-operative) data allowing input dataset dimension reduction

In our hybrid input dataset, each patient was assigned an input vector of measured or simulated biomarkers potentially available before operation. The hybrid dataset contained features from different semantic blocks: clinical data of different modalities (anthropomorphic, clinical assessment/diagnosis, instrumental data from ECG, echocardiography), CT/MRI derived indices, and simulations on a personalized ventricular model in both LBBB activation and BiV mode.

First of all, we excluded all mechanical dyssynchrony indices from the hybrid dataset as there were missed values in these features. Thus, we considered 31 features per patient for further analysis. To decrease the dimension of the input vector of biomarkers, we first analyzed pair-wise correlations between the features and found pairs having high correlation coefficients (r>0.75, p<0.05, see the hit-map in Figure 7 in Supplementary Materials showing color-scaled r-values between the features). While several input features demonstrated high correlations, we excluded one feature from every pair with r>0.85. This is a conventional practice in data analysis preventing information loss caused by a possible overestimation of the correlations between features due to the sample size being rather small.

We found no high correlations between the parameteres from the data-blocks of different semantics and modalities, suggesting that data from each block should be further used for CRT response classifier development. The only exception was a high correlation between clinical QRSd*_LBBB_* recorded before CRT operation and model-derived QRSd*_LBBB_* in the LBBB activation mode (r= 0.84). This correlation is not surprising because personalized LBBB models were tailored using ECG data recorded during electrocardiographic imaging using the Amycard EPI-system in patients in the LBBB activation mode with ventricular pacing turned off(see Methods Sec. for details). However, as this correlation did not overcome the threshold for data exclusion, both features were further used for classifier development.

Among the pre-operative clinical data, high positive correlations were found between LV EDV and ESV and between the corresponding linear LV dimensions EDD and ESD derived from echocardiography records before operation (r=0.97 and 0.94, respectively). The latter suggests the possibility of using either one dimensional biomarker as representative of both LV dimensions. Therefore, ESV and ESD features in each pair were excluded from the hybrid dataset used to develop a classifier.

Among the simulated features, we found high cross-correlations between the TAT/MTV and QRSd/MTV values (normalized to the survived myocardial tissue volume (MTV)) between each other in the same mode of activation and between each of them in the LBBB and BiV pacing (see numbers in Fig. 7). However, only QRSd/MTV*_LBBB_* and QRSd/MTV*_BiV_* showed a high correlation (r=0.90), which allowed to exclude the former feature from the dataset for classifier building. We found also some high correlations between simulated electrical dyssynchrony parameters and TAT under LBBB, and between the two inter-LV dyssynchrony indices under BiV pacing, but no one pair overcame the threshold for data exclusion (see Fig. 7).

Thus, we excluded 3 input features from the dataset. Finally, the hybrid dataset containing 27 input features per each of 57 patients was used further for classifier development (see the complete feature list in Figure 4).

**Figure 4:**
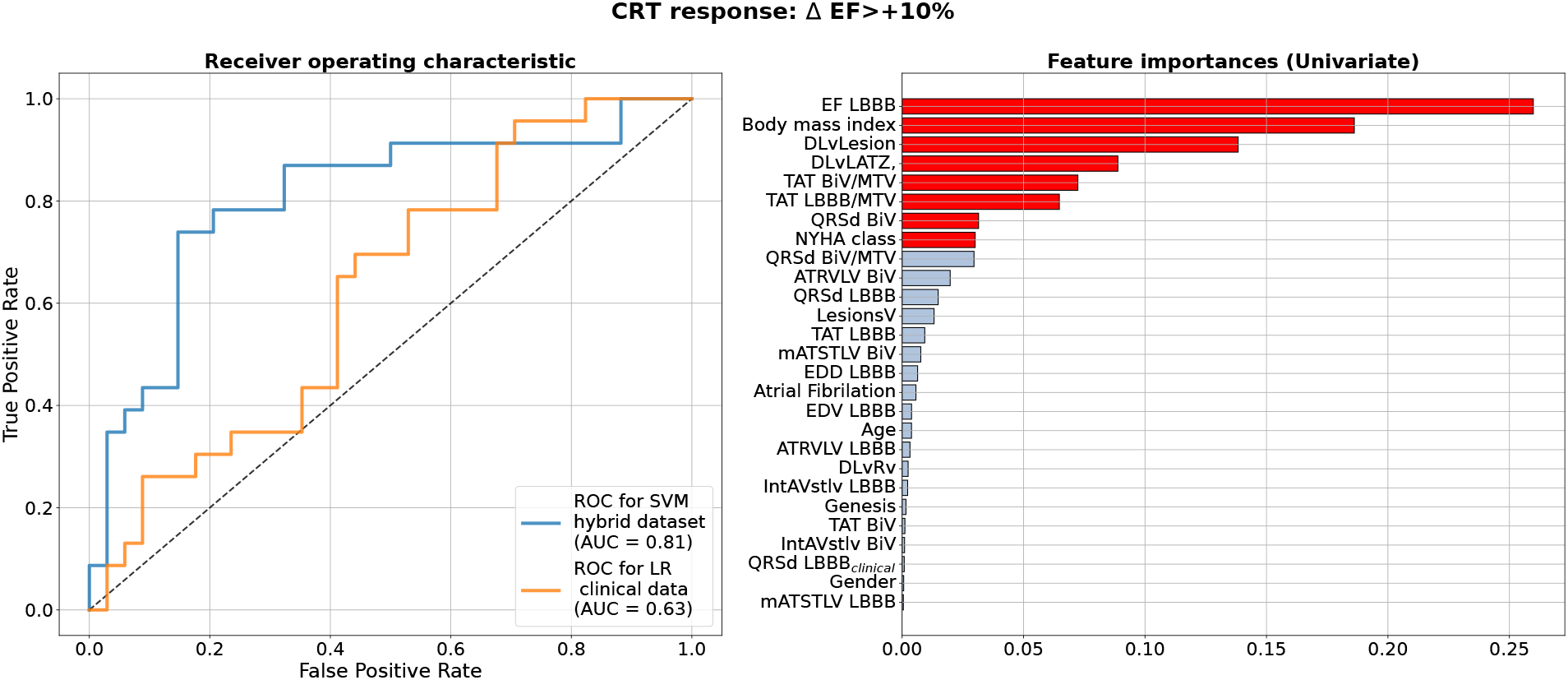
Best Machine Learning Classifiers for CRT response prediction from the hybrid dataset of clinical and model-drived data for 57 patients. **Left panel** shows receiver operating characteristic (ROC) curves for the best classifiers based on the ΔEF>10% criterion of CRT response. Blue line shows ROC curve for Support Vector Machine Classifier (SVM) using Leave-One-Out cross-validation on hybrid dataset. Yellow line shows a ROC curve with corresponding ROC AUC for a Logistic Regression(LR) model trained on the data subset containing clinical features as suggested in [13]. Values of the area under the ROC curve (ROC AUC) for the models are shown on the panel. **Right panel** shows clinical and model-drived feature list in descending order of importance ranged using Univariate feature selection approach for the best classifier.

### 3.3 Correlations within the output (post-operative) data

We performed a similar pairwise cross-correlation analysis for post-operative clinical features derived from ECG and echocardiography recordings, which are conventionally used to characterise CRT response in patients (see Figure 8 in Supplementary Materials). The CRT outcome dataset contains QRSd, EDV, ESV, and EF a year after CRT device implantation and their relative change against pre-operative values. High correlations (r>0.75) were found between EDV*_CRT_* and ESV*_CRT_*, and between ΔEDV*_CRT_* and ΔESV*_CRT_*. These findings support stronlgy the criteria we chose to identify CRT responders in our patient cohort using a separate analysis of either ΔEF*_CRT_* or ΔESV*_CRT_*, as these showed moderate dependence on each other (r=-0.61) and low-to-moderate dependence on other output features (Fig. 8).

### 3.4 Correlations between input and output features

Before building CRT response classifiers using ML approaches we analysed pairwise cross-correlations between input features and post-operative CRT outputs (see the correlation hit-map in Figure 9 in the Supplementary Materials). Our aim was to find potentially most predictive features from our input hybrid dataset and compare the feature importances assessed by linear regression with those selected by ML classifiers. The highest correlation (r=0.81) is seen between simulated QRSd*_BiV_* in BiV mode and clinical post-operational QRSd*_CRT_*. The correspondence between the simulated and clinical features reflects a good quality of personalized models fitted to post-operational ECG data recorded during BiV pacing by means of electrocardiographic imaging (Amycard system), and justifies the use of model-driven features in BIV mode for CRT response prediction. A high correlation was also found between clinical QRSd*_LBBB_* before operation and ΔQRSd*_CRT_* (r=-0.74) suggesting higher electrical synchronization in patients with initially shorter QRSd.

Other input-output features show either low or moderate correlations (see Fig. 9). In particular, post-operational ΔEF*_CRT_*, which we used to define CRT response, correlates with clinical BMI and pre-operational EF*_LBBB_*, and some model-derived features: distance from the LV pacing site to the late activation zone, normalized QRSd/MTV*_LBBB_*, TAT/MTV*_BiV_* and QRSd*_BiV_*. At the same time, ΔESV*_CRT_* showed correlations only with EDV*_LBBB_* and ESV*_LBBB_* and with the distance from LV pacing site to the area of LAT. For some input features we found no correlations with any of the output indicators, e.g. for simulated LBBB indices of ventricular electrical dyssynchrony.

Summarising the results of our analysis, we found a variety of relationships between clinical data before and after CRT, as well as between model-driven indices and clinical indicators of response. We hypothesise that combining clinical and simulated features can significantly improve prognostic models of the CRT response.

### 3.5 Predictive models of CRT response built on hybrid dataset of clinical data before operation and personalized model simulations of biventricular pacing outcomes

We used the hybrid input dataset containing 57 data entries with features derived from clinical data recorded prior to operation, CT/MRT derived data and simulated features calculated using personalized models of ventricular excitation in LBBB and BiV pacing activation modes for every patient from our cohort as described in the previous sections. We trained supervised classifiers using an EF10 criterion (ΔEF>10%) of CRT response. To choose the best classifier, we compared 4 different classification models (classifiers) with Leave-One-Out and five-fold cross-validation and 3 different feature selection methods inside a cross-validation loop. A summary of the model ROC AUC used to characterise the quality of the trained models is shown in Table 4. It is seen that average ROC AUC vary from a smallest value of 0.7 to the best one of 0.82 obtained for SVM and LDA classifiers with Univariate approach for feature selection.

**Table 4:** Comparison of ROC AUC for different Machine Learning classifiers using leave-one-out and five-fold cross-validation and different feature selection algorithms for EF10 criterion of CRT response.

| Classifier | Leave-one-out |  |  | Five-fold |  |  |
| --- | --- | --- | --- | --- | --- | --- |
|  | Feature selection method |  |  | Feature selection method |  |  |
|  | L1 | MDA | Univariate | L1 | MDA | Univariate |
| Logistic Regression | 0.76 | 0.78 | 0.80 | $0.74 \pm 0.15$ | $0.76 \pm 0.14$ | $0.80 \pm 0.14$ |
| Linear Discriminant analysis | 0.73 | 0.76 | 0.82 | $0.71 \pm 0.15$ | $0.76 \pm 0.15$ | $0.80 \pm 0.14$ |
| Support Vector Machine | 0.70 | 0.73 | **0.82** | $0.72 \pm 0.15$ | $0.75 \pm 0.15$ | **$0.80 \pm 0.14$** |
| Random Forest | 0.73 | 0.72 | 0.73 | $0.72 \pm 0.16$ | $0.70 \pm 0.16$ | $0.72 \pm 0.16$ |

L1 - Logistic Regression feature selection;
MDA - Mean Decrease Accuracy;
Univariate - Univariate statistical testing: two-sample t-test for continuous variables and chi-squared test for categorical variables;

Figure 4 (left) shows a ROC curve for the best SVM classifier trained for the EF10 response criterion. Table 5 summarises the classifier characteristics. The best SVM classifier for CRT response demonstrates a high accuracy of 0.82, sensitivity of 0.85, and specificity of 0.78.

**Table 5:** Performance of the classifiers on hybrid versus clinical data

| SVM on hybrid dataset |  |  |  |  |
| --- | --- | --- | --- | --- |
| $\Delta EF > +10\%$, cutoff = 0.46 | | | | |
|  |  | Responder | Non-responder |  |
| Model Responder |  | 18 | 5 |  |
| Model Non-responder |  | 5 | 29 |  |
| Accuracy | Sensitivity | Specificity | ppv | npv |
| 0.82 | 0.85 | 0.78 | 0.78 | 0.85 |
| LR on hybrid dataset |  |  |  |  |
| $\Delta EF > +10\%$, cutoff = 0.46 | | | | |
|  |  | Responder | Non-responder |  |
| Model Responder |  | 15 | 8 |  |
| Model Non-responder |  | 7 | 27 |  |
| Accuracy | Sensitivity | Specificity | ppv | npv |
| 0.74 | 0.65 | 0.79 | 0.68 | 0.77 |
| LR on clinical dataset |  |  |  |  |
| $\Delta EF > +10\%$, cutoff = 0.51 | | | | |
|  |  | Responder | Non-responder |  |
| Model Responder |  | 8 | 12 |  |
| Model Non-responder |  | 15 | 22 |  |
| Accuracy | Sensitivity | Specificity | ppv | npv |
| 0.53 | 0.65 | 0.35 | 0.40 | 0.59 |
| Response score by Feeny et al.[13] |  |  |  |  |
| $\Delta EF > +10\%$, cutoff = 0.62 | | | | |
|  |  | Responder | Non-responder |  |
| Model Responder |  | 13 | 15 |  |
| Model Non-responder |  | 10 | 19 |  |
| Accuracy | Sensitivity | Specificity | ppv | npv |
| 0.56 | 0.56 | 0.57 | 0.46 | 0.65 |

SVM - Support Vector Machine;
LR - Logistic Regression;
ppv - positive predictive value;
npv - negative predictive value;

ML scores generated by the best SVM classifier correlate with post-operational improvement in the EF + (r=0.46, p<0.001, see Fig. 5). Moreover, the distributions of the average scores in the responder and nonresponder groups in our patient cohort significantly differ between each other with a significantly higher average score in the responder versus nonresponder group (0.58±0.25 *vs* 0.29±0.22, p<0,01, see Table 3, Fig. 6). A corresponding score threshold of 0.46 was defined for the responders in our patient cohort for the best classifier according to the EF10 response definition.

**Figure 5:**
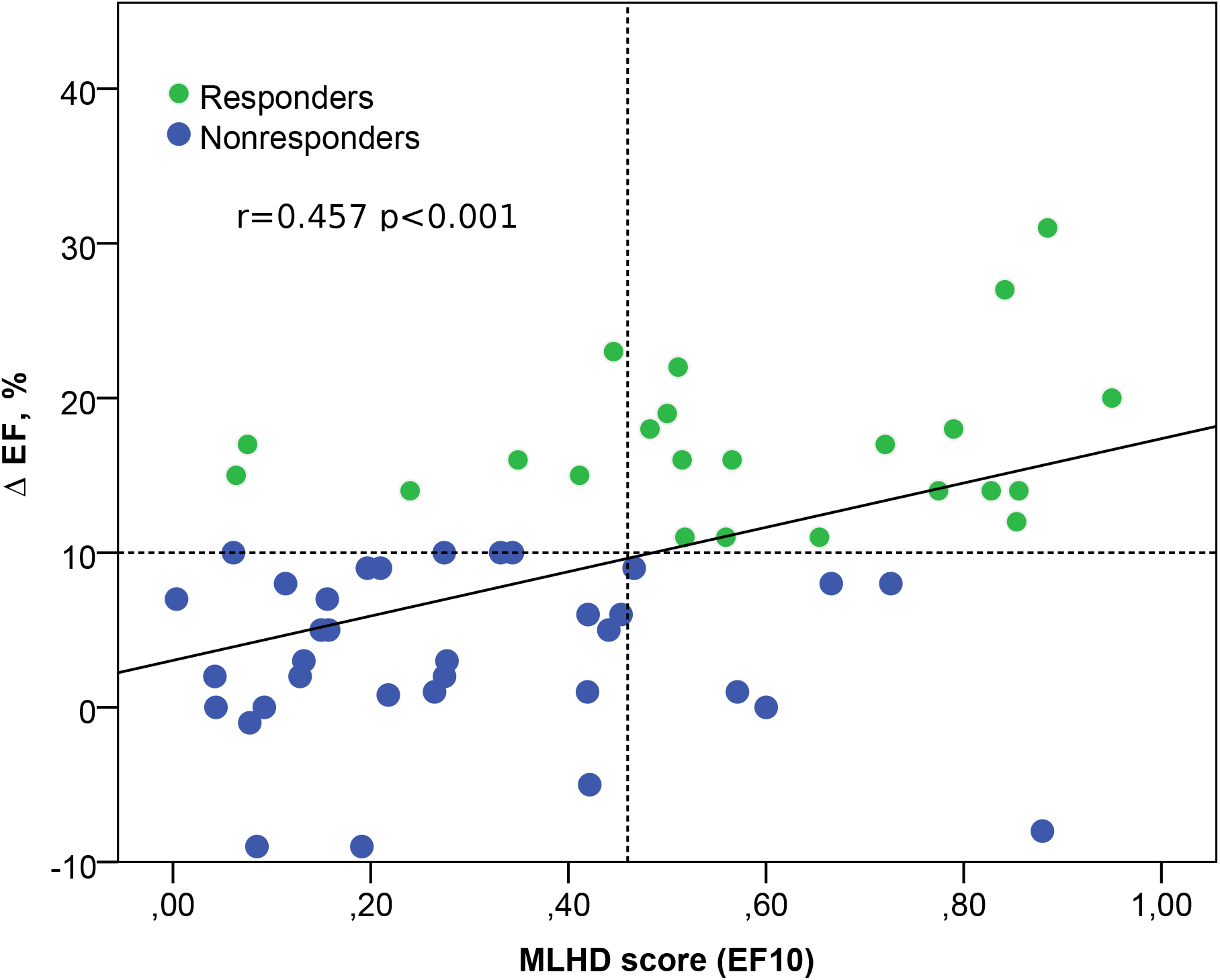
Relation between the ML score on the hybrid data (MLHD score) for EF10 criterion and the post-operational change in the EF. Solid line - regression line Δ EF = 3 + 14 MLHD score; horizontal dotted line shows a 10% threshold for LV EF improvement; vertical dotted line is a MLHD score cutoff of 0.46 for responders; r is the Spearman correlation coefficient; p is the significance for the group difference.

**Figure 6:**
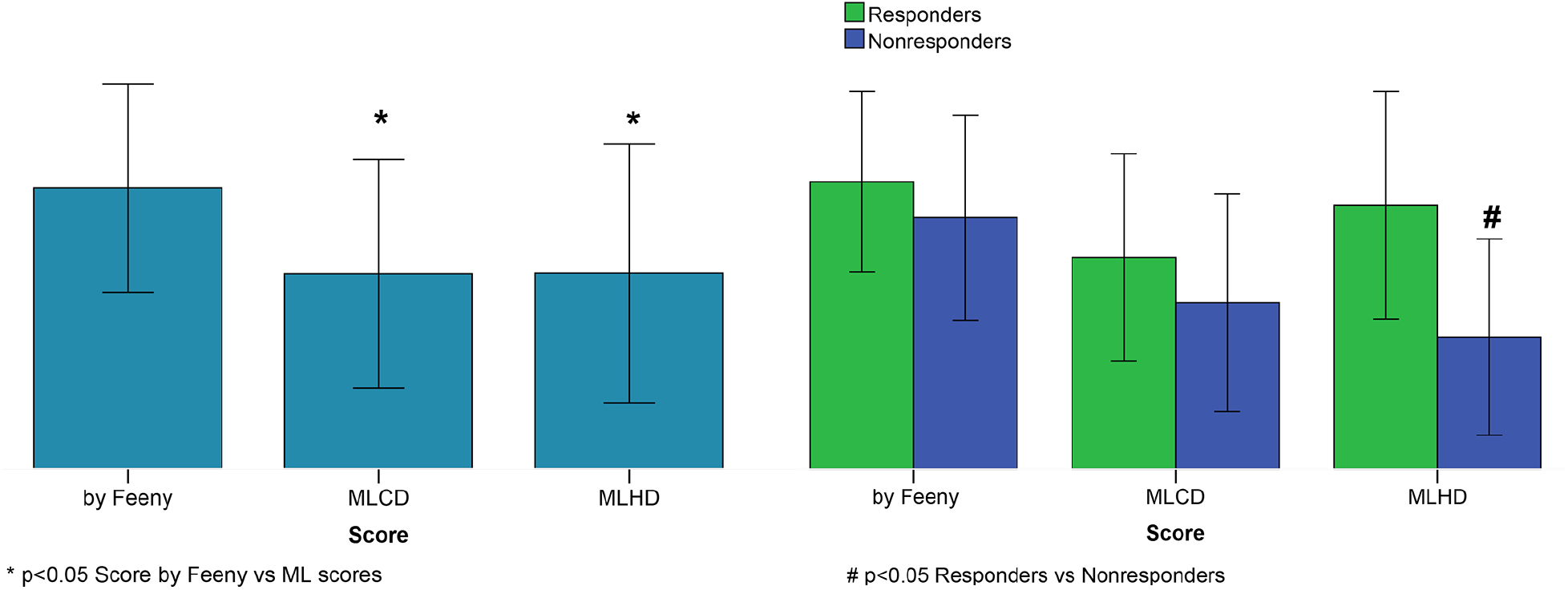
CRT response scores. Left panel: Average scores. Right panel: Average scores for responders and nonresponders. Score by Feeny et al [13]; MLCD – ML score on the clinical data for EF10 criterion; MLHD - ML score on the hybrid data for EF10 criterion. Bar indicates mean. Error bar is SD. Nonparametric Friedman’s two-way ANOVA was used to compare related groups (Score by Feeny vs ML scores). Comparison between two independent groups (responders vs nonresponders) was performed using the Mann-Whitney test.

Figure 4 (right) shows a ranged list of feature importances selected by the SVM classifier trained on the entire dataset for the EF10 CRT response criterion. Eight most important features were selected for the final classifier. The pre-operative EF*_LBBB_* showed the highest importance among other inputs, which is in line with our findings on the correlation between ΔEF*_CRT_* and EF*_LBBB_*. The other two of the three clinical features contributing to the CRT response were BMI and NYHA stage. Therefore, the majority of the selected features were indices derived from CT/MRI and simulated features in the LBBB and BiV modes of activation. In particular, the distance between the LV pacing site and the lesion zone was the third in the feature importance range, and a combination of TAT/MTV*_LBBB_*, TAT/MTV*_BiV_* and QRSd*_BiV_* showed the highest importance among simulated features. Corresponding coefficients at the input variables in the terms of the best LR classifier are given in Table 6 in the Supplementary Materials.

The yellow line in Figure 4 (left panel) shows the ROC curve for an LR classifier trained on the clinical data only according to the EF10 criterion. The sub-set of clinical features used here for CRT response prediction was the same as selected in the article by Feeny and co-authors [13] for their best LR classifier (see the complete feature list and corresponding coefficients at the input variables in Table 6 in the Supplementary Materials). The average ROC AUC for this predictive model appears to be 0.63 for our patient cohort, with an average accuracy of 0.53, sensitivity of 0.65, and specificity of 0.35 (Table 5), which are much lower than the characteristics of the ML model trained on the hybrid input dataset containing a combination of clinical and model-driven features. Note that this AUC is close to the AUC value of 0.62 we obtained for the LR classifier trained on EF*_LBBB_* only, suggesting that the rest of the clinical information does not contribute essentially to the model predictions.

We compared also the accuracy of EF10 improvement predictions from our ML classifier on the hybrid data with the accuracy of predictions based on the ML scores calculated using a calculator provided in the Supplementary Material in [13] (Table 5). Predictions proposed by the ML score calculator on our patient data showed an accuracy of 0.56, sensitivity of 0.56, and specificity of 0.57, which are similar with the performance of the LR classifier trained on the clinical data, but much lower that the performance of our ML classifiers on hybrid data.

Figure 6 shows average ML scores generated by the classifier on the hybrid data for the entire patient cohort and for the responder and nonresponder groups according to the EF10 definition in comparison with the ML scores predicted on the clinical data only and those from the calculator by Feeny et al. [13]. The average ML score from [13] on the entire patient cohort is seen to be higher than our ML scores, explaining lower rates of true positive and true negative predictions from the calculator on our patient cohort. Moreover, the only classifier on hybrid data generates significantly higher ML scores in the responder versus nonresponder group, suggesting its higher predictive performance. In contrast, the average ML scores did not differ between responders and nonresponders according for the LR classifier on the clinical data (0.47±0.23 *vs* 0.37±0.24, p=0.111) and for the calculator from [13] (0.63±0.20 *vs* 0.55±0.23, p=0.213) in our patient cohort (see also Table 3).

These results clearly highlight the significance of model-driven features for CRT response prediction.

## 4 Discussion

The researchers sought ways to predict CRT response for ensuring more effective patient stratification and different outcome end-points for improving state, increasing survival period and preventing adverse effects [33]. Despite of intensive research performed in the field, the fraction of patients with low response to the therapy remains as high as 30-50% depending on which criteria are used for assessing CRT outcome. New artificial intellegence and ML based approaches to data analysis have been extensively used in attempts to increase the accuracy of patient differentiation [7, 9, 13, 12]. Computational models based on clinical data are also employed to identify mechanisms responsible for the poor efficacy and develop approaches improving CRT outcomes [14, 34, 35, 36]. Recently, a new trend has emerged in this research area, which uses a combination of clinical and model data together with ML for solving challenging medical problems [37, 38]. As far as we know, there have been no reports of in-silico studies involving a hybrid approach to predict CRT response in a cohort that would combine a dataset of patient-specific features derived from clinical measurements and simulations on personalized ventricular models.

### 4.1 Improvement of classification models built on hybrid data versus predictors on clinical data

In this study, we combined the MR/CT-imaging and model derived features with pre-operative clinical data used conventionally to characterise patient’s state in a hybrid dataset for building predictive models of CRT response in the patients by ML techniques. A sub-set of simulated features containing TAT, QRSd and three electrical dyssynchrony indices generated by every of 57 patient-specific electrophysiology models under LBBB and BIV pacing was used as an input to ML algorithms. The personalized models were also used to define LAT zone in the LV under LBBB mode of activation and to calculate the distances between the pacing sites, and from the LV pacing site to the LAT zone and to the LV lesion zone, which were also used as input features for ML classifiers. The basic hypothesis of our study was that model-driven simulations of the response to BiV pacing may essentially enhance the predictive power of the hybrid dataset for CRT response evaluation.

Despite the rather small size of the dataset used for ML classifier development (57 entries in the entire dataset), we were able to obtain ML classification models achieving high accuracy in predicting the response to CRT (see Table 4 and Figure 4). The ROC AUC value for the best SVM classifier is as high as 0.82 for ΔEF>10% cutoff for responders.

The most significant result of our study is that our best classification models built on the hybrid dataset show notably higher accuracy as compared with the ML classifier trained on the pre-operative clinical data only (see Fig. 4, Table 5). In the latter, we used a data sub-set containing 8 clinical features selected for the best LR model suggested by Feeny and co-authors in their recent study [13] and trained a model for our cohort. The best model built on the clinical data demonstrated a ROC AUC of 0.63, and the accuracy, sensitivity and specificity much lower than those for the classifiers built on the hybrid dataset (see Fig. 4 and Table 5). Predictions proposed by the ML score calculator provided in the Supplementary materials of the article [13] and tested on our patient data also showed a low accuracy of 0.56, which is similar with the performance of the LR classifier trained on the clinical data for our patient cohort. Comparing the characteristics of different classifiers on clinical data from about thousand of patients reported in [13] (see Table 3 ibidem), we may conclude that our ML classifiers built on the combination of clinical and model-derived features demonstrate significantly improve prediction quality with higher accuracy, sensitivity and specificity.

In addition, we compared the average ML CRT response scores in the responder and non-responder groups provided by the best SVM classifier on hybrid data, the LR classifier on the clinical data and that provided by the ML score calculator from [13] (see Fig. 6). Noteworthy, for the SVM classifier based on the hybrid data, we found a significantly higher average score in the responders versus nonresponders confirming the predictive power of the ML model. In contrast, the average ML scores predicted by the LR classifier on the clinical data and calculator from [13] did not differ between responders and nonresponders in our patient cohort (see Table 3).

Therefore, our results clearly show significant advantages ensured by the use of hybrid data combining clinical data with simulated features from personalised electrophysiology models for building ML predictive models of CRT response.

### 4.2 Feature selection for classification models from hybrid data

During classifier development, we tested several feature selection methods for different classifiers and different numbers of features to define the final model with best characteristics (see Fig. 4). Note that we did not predetermine input features for classifiers based on prior analysis. Instead, the features were automatically selected inside the cross-validation loop as described in Methods section. The final feature lists selected for the best predictive models contain 8 inputs. Importantly, the most important feature set contained fewer clinical features compared with model-derived ones. In consistency with ESC guidelines on the significance of pre-operative baseline LV EF*_LBBB_* for CRT response, it was selected as the most important feature for the classification model based on the EF10 definition (see Fig. 4). Interestingly, BMI was selected at the second position in the feature chart. The latter result is in line with study by Hsu and co-authors [39], who demonstrated that BMI<30 kg/m^2^ predicted LV EF super-response.

We also tested the importance of model-driven characteristics extracted from the CT/MRI data coupled with model simulations. In our study, LV lesion volume (both absolute and relative to the survival myocardium volume) did not reveal high importance by itself, but the distance from the LV pacing site to the lesion zone was selected as the third most important feature for classifiers (see Fig. 4). We found no significant correlations between the distance from LV pacing site to the lesion zone and the post-operative values of LV EF improvement ΔEF*_CRT_* or ESV reduction ΔESV*_CRT_* (see Fig. 9 in the Supplementary materials). However, the role of the distance from the LV pacing site to the lesion in CRT response prediction was supported by a positive correlation between the ML score and the distance (r=0.445, p=0.001) and much higher average distance in the responder versus nonresponder group (45±28 *vs* 28±27 mm, p=0.02, see Table 3).

Our findings are consistent with the results of clinical studies which assessed the significance of myocardial lesion size for CRT response. The extent of scar core and gray zone was automatically quantified using cardiac MRI analysis [40]. The highest percentage of CRT response was observed in patients with low focal scar values and high QRS area before operation. Such area was calculated using vector-cardiography. In study by Marsan and co-authors [41] MRI was performed in candidates to derive LV mechanical dyssynchrony and the extent of scar tissue to predict CRT response. Higher LV dyssynchronies were strongly associated with echocardiographic response to CRT, while the total extent of scar correlates with non-response. Importantly, a univariable logistic regression analysis showed that the presence of a match between the LV lead position and a transmural scar was also significantly associated with non-response to CRT. The location of scar in the posterolateral region of the LV, which is empirically thought to be a target site for LV lead implantation, was associated with lower response rates following CRT [42]. In study by Pezel and co-authors [43], no difference was found in presence and extent of scar between CRT responders and non-responders. However, in non-responders, the LV lead was more often over an akinetic/dyskinetic area suggesting the presence of tissue lesions, a fibrotic area, or an area with myocardial thickness < 6 mm.

As seen in Figure 4, the distance from the LV pacing site to LAT zone was selected as a 4-th feature in the importance list for EF10 definition of CRT response. Accordingly, we revealed a low negative correlation between the distance and ML classification score (r=-0.263, p=0.048) suggesting its possible role in CRT response prediction. This was a bit surprising, as no difference in this feature was found between the responders and non-responders (see Table 3), as well as no correlation with LV EF improvement in our patient cohort (r<0.25, p>0.05). However, selection of this distance as an important feature for ML classifier is in line with clinical studies, where the LAT zone was considered as a target area for LV lead deployment [44, 33, 45, 46]. In particular, consistent with clinical data, our results indicate that optimal electrode deployment should be guided by a kind of minimum-maximum optimization with respect to the distance from LAT and lesion areas, respectively. Preoperative model-based prediction of such optimal pacing site location seems extremely valuable.

Although ventricular mechanical dyssynchrony was considered with respect to CRT improvements [47, 48, 46, 49], we did not use mechanical dyssynchrony indices in developing our classifiers because not every patient had these features indicated in the retrospective dataset. We did not find a correlation between the ML response scores generated from the selected hybrid data and the mechanical dyssynchrony indices measured in 34 patients at the baseline (r<0.25, p>0.05 for IVD, Tsmax-Tsmin, SD12). This was not consistent with a correlation between the IVD index and postoperative ΔEF in the patient cohort (r=0.32, p=0.029), and a significant difference in the average IDV indices between responders and nonresponders defined by LV EF improvement (75±17 *vs* 63±19, p=0.013, see Table 3). These controversial findings did not allow us to disprove the possible importance of mechanical dyssynchrony indices for ML response prediction, and this hypothesis should be further evaluated on a dataset of bigger size.

It is especially remarkable that each classification model included simulated characteristics of myocardial activation and ECG from the personalized electrophysiology models under LBBB and BiV pacing selected among the most important features. Our best SVM classifier for the EF10 response definition selected three simulated features TAT/MTV under LBBB and BiV pacing, and QRSd under BIV pacing among the 8 most important ones for EF improvement prediction (see Fig. 4). In particular, two of the three features TAT/MTV and QRSd under BIV pacing correlated with EF improvement (r=0.27 and r=-0.31, p<0.05, see Fig. 9 in Supplementary Materials), supporting their importance for ML predictions. Note, the *in-silico* indices of electrical dyssynchrony assessed in our study were not selected as important for ML classifiers. These indices were previously suggested by Villongco and co-authors [20], who demonstrated a correlation between the post-operational ESV reduction and the change in the mAT*_ST LV_* index of inter-ventricular dyssynchrony under BiV pacing against the LBBB baseline on data from 8 patients. In study by Lumens and co-authors [36], a combination of clinical data and personalized models of cardiac mechanics and hemodynamics also demonstrated significant role of inter-ventricular electrical dissynchrony in predicting CRT response defined by an improved LV hemodynamic performance assessed via increase in the maximal derivative of LV pressure (dP/dtmax). In contrast, we found no significant correlations between any of the simulated indices of electrical dyssynchrony and echocardiographic CRT response in our cohort (r<0.25, p>0.05). The role of such simulated indices needs further analysis to be performed on a dataset of bigger size.

### 4.3 Hybrid dataset size and cross-validation

It is noteworthy that the ML classifiers we developed to predict LV EF improvement can be considered as powerful, especially taking into account the database size of less than 60 entries. In several studies, the ROC AUC was shown to improve significantly with increasing the dataset size from tens to thousands of entries [13]. These results allow us to expect further substantial improvement of the quality of the ML classifiers with further increasing the training dataset size. Poor reproducibility of ML results is known as a frequent problem with classifiers developed on small samples. In our case, the restrictive size of the dataset did not allow us to divide data into a conventional 80% training sub-set and 20% testing sub-set, so we had to use 57 Leave-One-Out combinations of data for classifier training.

To confirm the good quality of our classifiers, we tested also a widely-used repeated stratified five-fold cross-validation method with over 1000 iterations. In this approach we chose 1000 combinations of 45 training samples from our dataset to train classifiers and the rest 12 samples to test the models. The statistics of the ROC AUC for the five-fold cross-validation approach is shown in Table 4 (right column) in comparison with that of Leave-One-Out cross-validation (left column). It is seen that average ROC AUCs at five-fold cross-validation are slightly lower than those generated with the Leave-One-Out approach but the latter values fall in the confidence interval of ROC AUC distributions shown by five-fold cross-validation on our dataset. Results demonstrate stability of the ML classifiers we built on our hybrid dataset and confirm the robustness of the ML predictions.

When developing our classifiers, we noticed that the list of features in the cross-validation loop was not steady. This was mainly due to the small sample size. However, even taking this factor into account, we obtained a high accuracy of the constructed classifiers. Testing the SVM classifier for EF10 with a smaller number of features, we found that even for 5 features, the classifier showed the same accuracy as for 8 features (for other criteria of CRT response we observed a lower accuracy with reduced input data dimension). This suggests that with any further increase in the dimension, the classifier cannot converge to optimal solution. Therefore, adding more features to such a small size dataset does not make classifiers more accurate. We hope that with an increase in the size of dataset, the accuracy of the classifiers will additionally increase due to a more stable feature selection.

### 4.4 Classifiers for various definitions of CRT response

We used different CRT response definitions to build ML classifiers for our hybrid dataset. Unfortunately, no consensus has been achieved on how to define ‘response’ to CRT [30], making it difficult to compare different clinical trials and modelling studies. CRT response definition by markers of LV reverse remodeling following device implantation is widely used, and a more than 15% reduction in LV end-systolic volume (ΔESV<-15%) is the most widely accepted criterion [31]. In consistency with that, an optimal cutoff value for ΔESV was defined at 13.5% (sensitivity=0.719, specificity=0.719) for a 1-Year hierarchical clinical composite end point in patients who underwent CRT [50]. Our earlier 278 patients’ study by Chumarnaya and co-authors [51] revealed a 9% cutoff value for ESV reduction for responders. Surprisingly, in our patient cohort, the grouping by either 10% or 15% cutoff for ESV reduction for responders was the same. Therefore, we used the latter definition (ESV15) to determine a positive response to CRT.

A summary of the statistics for the hybrid dataset labelled according to the ESV15 definition of CRT response is presented in Table 7 and described in the Supplementary Materials. ML classifiers with leave-one-out cross-validation on the hybrid data showed a high performance with best ROC AUC of 0.74 (see Fig. 11), and an accuracy of 0.70, sensitivity of 0.87, specificity of 0.37, ppv of 0.73 and npv of 0.58. See also the classifier characteristics for five-fold cross-validation in Table 9. The ML scores generated by the best classifier built on the ESV15 criterion correlated with post-operational reduction in ESV (r=-0.27, p=0.039, see Fig. 12).

The results are slightly less powerful as compared with classifiers built on the EF10 criterion. The latter showed higher ROC AUCs, similar sensitivity, but higher specificity as compared to ESV15 (see Figs. 4 and 11, and Tables 4 and 9). The ML scores based on ESV15 labelling are higher as compared with EF10 scores (0.69±0.18 *vs* 0.40±0.35, p<0.01, respectively) tending to overestimate predictions for the negative response. Note that for both CRT response criteria the average scores are significantly higher in responders *vs* nonresponders, indicating good predictive quality of the ML classifiers.

Surprisingly, the sub-sets of 8 most important features selected for classifiers on different response criteria almost did not intersect. For the ESV15 criterion, the pre-operative EDV*_LBBB_* showed the primary importance among other inputs in consistency with its correlation with ΔESV*_CRT_* (r=-0.36, p<0.05). Another clinical feature selected for classification was IHD/DCM index reflecting the aetiology of CHF in patients (see Fig. 11 in Supplementary materials). The rest of the selected features were indices derived from CT/MRI data and simulated features in LBBB and BiV modes of activation. Similar to EF10, the distance from the LV pacing site to lesion zone was the third in the feature importance range, and simulated TAT/MTV*_LBBB_* was selected for both ESV15 and EF10 criteria together with other model-derived features different between the criteria (see Figs.4 and 11).

Unexpectedly, we were not able to generate a predictive model for ESV15 criterion from the clinical feature sub-set suggested in [13] with ROC AUC>0.5 on the dataset for our patient cohort. When we calculated ML scores using the calculator from [13] for our responders and nonresponders defined by ESV15 criterion, the average ML scores did not differ between the groups, while the ML scores based on the hybrid data were significantly different (see Table 7). These findings also point to the power of model-driven data in CRT response prediction.

We also compared the accuracy of ML classifiers built on the hybrid dataset for CRT response defined by 5, 10, 15% LV EF improvement and by coupled EF10 and ESV15 criteria (see Tables 8, 9 in the Supplementary Materials). For every response definition, our best classifiers demonstrate improved performance as compared with all clinical and ML predictors reported in [13]. Note again that our hybrid data classifiers were trained on a dataset of much smaller size than previously published [13]. Like in [13], for different ΔEF criteria an average accuracy of the predictive models increases with the cutoff for the LV EF improvement for CRT responders. However, the sensitivity and predictive positive value of the models tend to decrease with increasing the ΔEF cutoff, while both the specificity and predictive negative value increase. Thus, ML scores tend to underestimate the probability of a super-response. The best balance between sensitivity and specificity was shown for the ΔLV EF>10% definition of CRT response which also demonstrates the best ROC AUC among other criteria, thus supporting the choice of this criterion for response evaluation in patients.

### 4.5 Principal coordinate analysis and unsupervised ML clustering for CRT response prediction

The classifiers we discussed in the previous sections were built using feature selection approaches where input values are original functional characteristics of the processes. Often, in ML algorithms principal component analysis (PCA) is used for data dimension reduction, which allows more objective exclusion of collinearity between the input features. We tested the PCA in combination with Logistic Regression using different numbers of PCs for classifier development. We evaluated ROC AUCs using from 2 to 10 PCs, and obtained the best ROC AUC of 0.70 for 5 PCs (Fig. 10, left panel, in the Supplementary Materials) with explainable variance of 0.58. This ROC AUC is much lower than the best values demonstrated by other ML classifiers we developed using row feature values.

As we showed in the previous section, classification results depend on the positive CRT response definition used for data labelling. Another ML approach is unsupervised ML data clustering based on their similarity without the help of class labels. We performed clustering by K-means, used in recent studies for CRT response evaluation [12, 11]. Using K-means clustering on the two first PCs, we differentiated our dataset into 2 clusters (see Figure 10, right panel, in the Supplementary materials). However, mean ΔEF*_CRT_* were not significantly different between the groups (8.0±8.6% *vs* 10.7±8.3%, p =0.27), similar to no difference in mean ΔESV*_CRT_* (-20±38% *vs* -34±31%, p= 0.15). Moreover, distribution between the two clusters of CRT responders and non-responders defined by the EF10 criterion shows random assignment to the groups (see Fig. 10, right panel, in the Supplementary Materials). These findings suggest that unsupervised learning on a small dataset does not allow one to reliably differentiate pre-operational data into groups clearly associated with CRT response characteristics. In contrast, the supervised ML algorithms we developed provided valuable predictions of CRT response showing the potential of model-derived features.

## 5 Limitations

There are several limitations in our study that have to be overcome to make our approach actually usable in clinic. First, ventricular geometry in our personalized models was derived from CT images obtained after CRT device implantation, not before it. This was essential for this proof-of-concept study because it allowed us to define the precise location of pacing electrodes and to fit our models to both LBBB and BiV ECG data for the same ventricular geometry thus demonstrating the potential of our models for reproducing real clinical data. Despite supposed difference in the ventricular geometry our simulated ECGs in the LBBB mode had a high correlation with pre-operative clinical ECGs (r=0.84, p<0.05), thus demonstrating the effect of ventricular geometry as being secondary. Of course, the reverse remodeling of the ventricles after CRT may affect the difference in model simulations before and after operation. That is why we primarily focused on the CRT response definition based on the EF improvement which has low-to-moderate correlation with ventricular remodeling in our patient cohort. The main idea of using model-derived biomarkers for CRT response prediction was the possibility to assess the primary effect of ventricular synchronization itself on the electrophysiological characteristics of activation, where changes in the geometry seem less important.

The second limitation is that we used a simplified approach to model personalization focusing on the mean QRSd from 12-lead ECG as a target for the parameter identification problem. Then we used the maximal QRSd as a model biomarker for building a classificator. The QRS morphology may provide much more information for tailoring personalized electrophysiology models and then for CRT response predictions. A recent study of Camps et al. [19] showed a way towards more accurate personalization of the activation processes in ventricles based on the QRS signals recorded in patients, which may be usefull for model improvement. Feeny et al. [12] also demonstrated the power of the entire ECG signal for ML predictions of CRT response, suggesting that the use of the entire simulated ECG under BiV pacing may further improve ML predictors.

Next, we have shown high importance of the distance from the LV pacing site to the lesion zone in ML predictions. In this study, we did not have access to raw MRI data from patients to be able to derive accurate information on post-infarction scar or fibrosis morphology. We used only textual descriptions of the infarct zone location with a segment accuracy within a 17-segment AHA LV model from an expert who evaluated MRI data in patients. There are great examples of using detailed morphology of myocardial lesions in personalized cardiac models for predicting the risk of cardiac arrhythmia and patient stratification [17]. We think more objective information on the scar and fibrosis morphology may improve predictive models of CRT response as well.

In this study, we used simulations from personalized ventricular electrophysiological models for CRT response prediction. The model choice is justified by the essence of the therapy, which ensures electrical synchronisation of ventricular activation, and the success of this synchronization determines the outcome of the operation. However, the goal of CRT implantation is the synchronisation of ventricular contraction and subsequent improvement in the mechanical performance of the ventricles. This opens up a further direction for studies using electromechanical models of cardiac activity which are being developed in modeling community including our group [34, 52, 53] and which are able to predict directly EF, dP/dtmax changes and other mechanical biomarkers of CRT response. Such models were already used for clinical data analysis in CRT patients by several groups [34, 54, 14, 55], demonstrating the power of such simulations for CRT response predictions. In particular, we believe that reduced mechanical models using regression or ML approaches to reproduce the behaviour of complex 3D models such as developed with our participation [56] would be the the best choice in terms of possible clinical application of model simulations.

The study also employed information on the precise location of pacing electrodes derived from post-operative CT imaging data. Correspondingly, the predictive models we developed used model-derived data in BiV pacing with pre-determined electrode location. If the models to be used for pre-operative CRT predictions, the pacing electrode location should be suggested using model simulations as well. A number of clinical and simulation studies paid great attention to the possibility to target pacing lead implantation [57, 58, 59]. Different criteria for optimal electrode location were discussed in the literature, including optimization of electrical synchronization characteristics, e.g. maximal narrowing of QRSd, or minimisation of interventricular dyssynchrony; maximal proximity to the late activation zone or late contraction zone; avoiding the match with infarct zone; or maximizing the mechanical performance characteristics, e.g. dP/dtmax [40, 58, 60, 34]. The use of simulations from personalized models in CRT response prediction opens the possibility to re-evaluate these hypotheses and suggest a new strategy for implantation planning with allowing for model-based prediction of optimal location for pacing electrodes. We are going to test this hypothesis in a future study.

The long-term goal of CRT is to reduce morbidity and mortality in heart failure patients with reduced left ventricular function and intraventricular conduction delay. Several studies tested ML approaches for predicting outcomes after CRT in terms of patient survival and frequency of adverse events in the longer term after operation [7, 8]. We had no sufficient data to perform such analysis using simulated data, and this could be another new direction of future studies.

Last but not least, in this study we had a limited data sample from 57 patients. Of course, this number is quite small for ML algorithms operating on thousands of entries with a possibility to use separate subsets for training and testing. However, our predictive models based on hybrid data from clinic and computational models of cardiac activity have demonstrated high performance with accuracy much higher than that demonstrated by predictors developed on clinical data from a thousand of patients. This inspires hope that a larger dataset and more informative data from time-dependent simulated signals may further improve CRT response predictions.

## 6 Conclusions

We have developed 57 personalized models of ventricular geometry and myocardial tissue lesions from post-infarction scars and non-ischemic fibrosis using MRI and CT imaging data recorded in patients who underwent CRT implantation. The models were used to simulate ventricular activation and ECG on the patient torso. Indices characterising total activation time, QRS duration at 12-lead ECG, and inter/intraventricular dyssynchrony in LBBB activation and BiV mode were then used for developing ML classifiers of CRT response on hybrid clinical and model-derived data.

Despite a limited dataset, we have developed several high-performing ML classifiers from the hybrid dataset. The best SVM classifier showed an accuracy of 0.82, sensitivity of 0.85, and specificity of 0.78, improving quality over that of ML predictors built from much larger datasets of clinical data in previous published studies.

The majority of the most relevant features selected from the hybrid dataset for the ML classifiers were model-driven indices, suggesting their great power for CRT response prediction. Distance from the LV pacing site to the scar zone and ventricular activation characteristics under BiV pacing were shown as the most important features among the selected model-driven indices for patient classification between responders and nonresponders.

## Funding

This work was supported by Russian Science Foundation grant No. 19-14-00134.

## Supplementary Data

### Correlation between features

Figures 7, 8, 9 summarized significant correlations found in the dataset.

**Figure 7:**
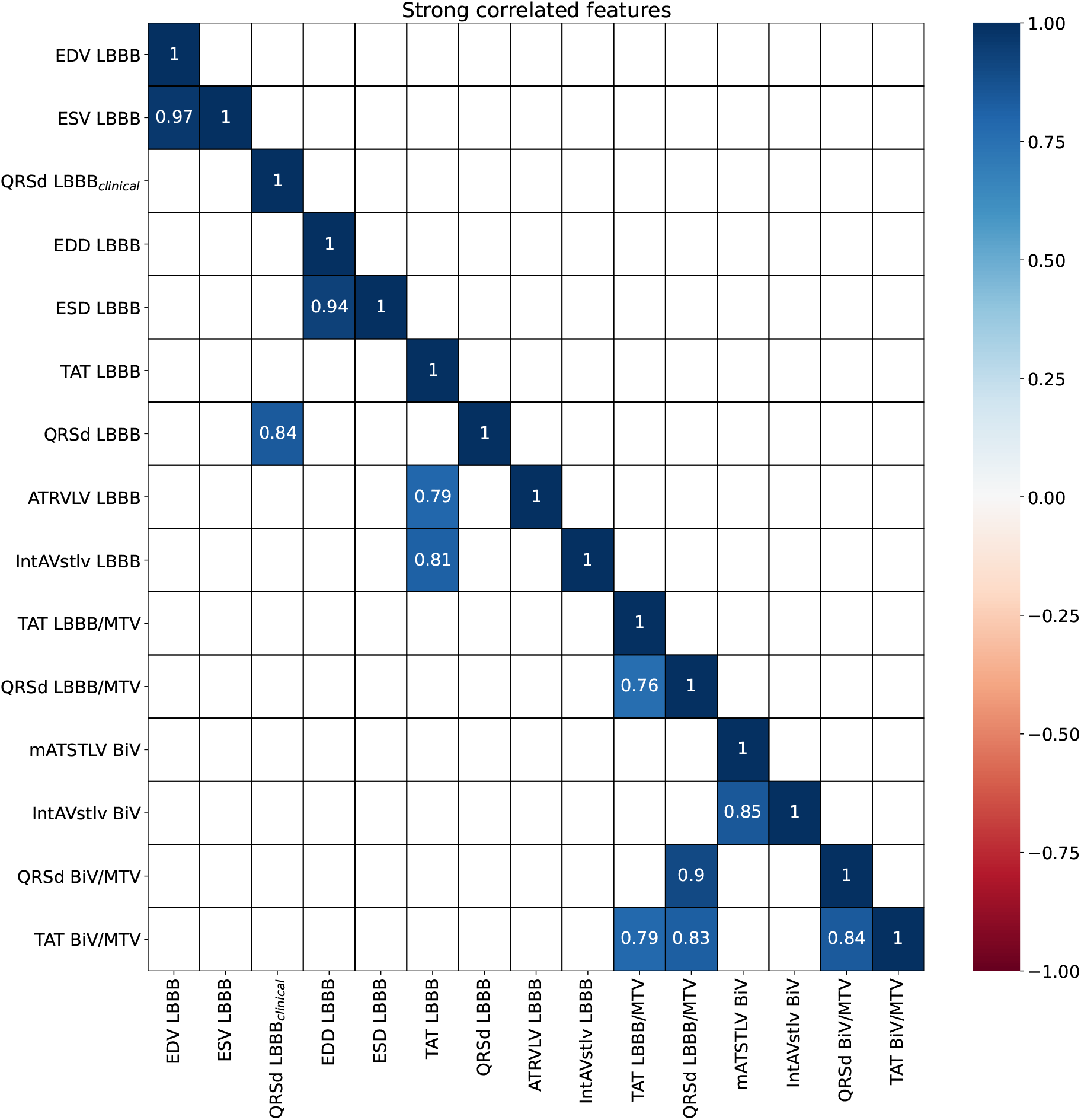
Correlation matrix of input features. Features with correlation coefficient r>0.75 and p<0.05 are shown.

**Figure 8:**
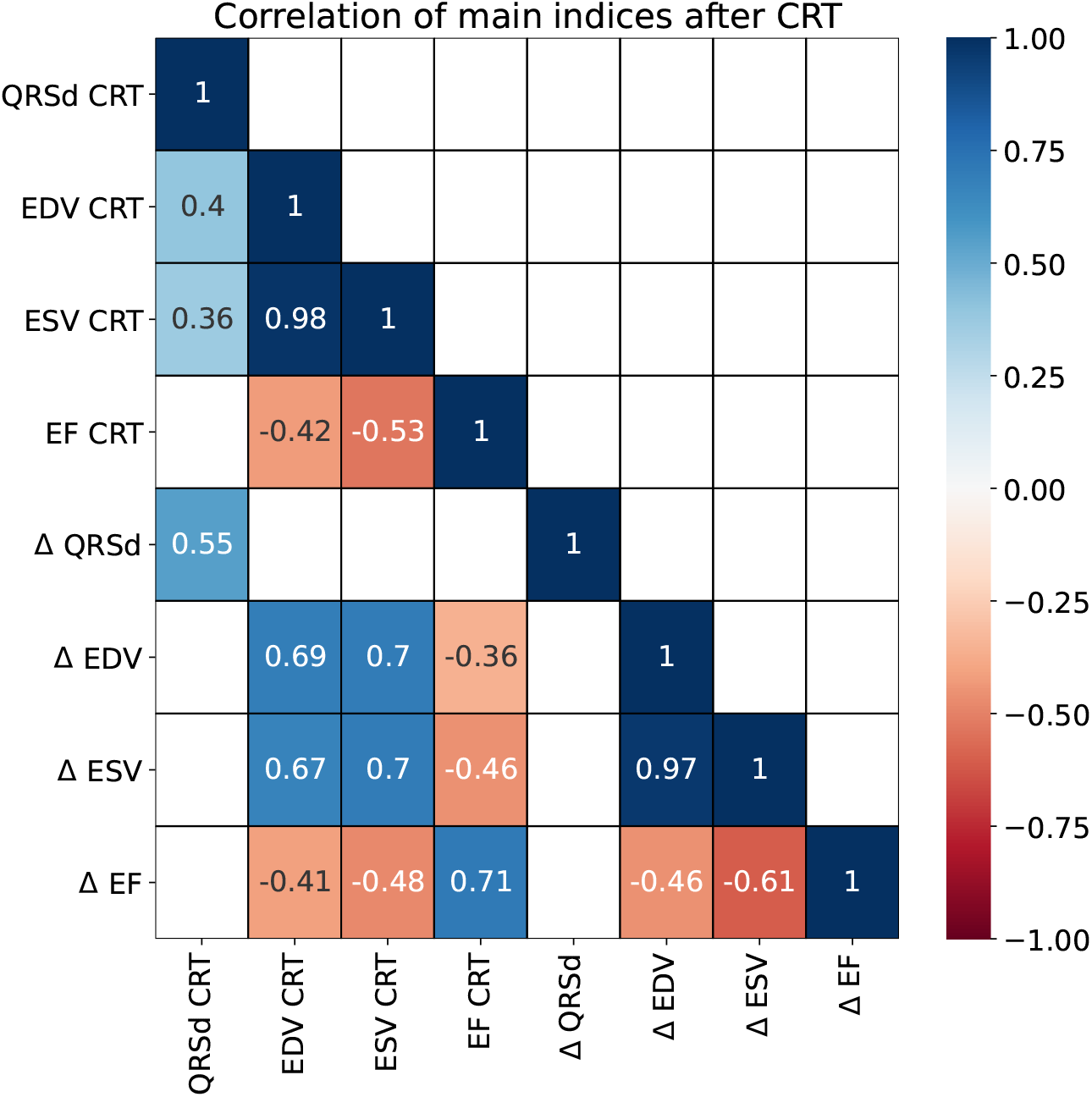
Correlation matrix of output features.

**Figure 9:**
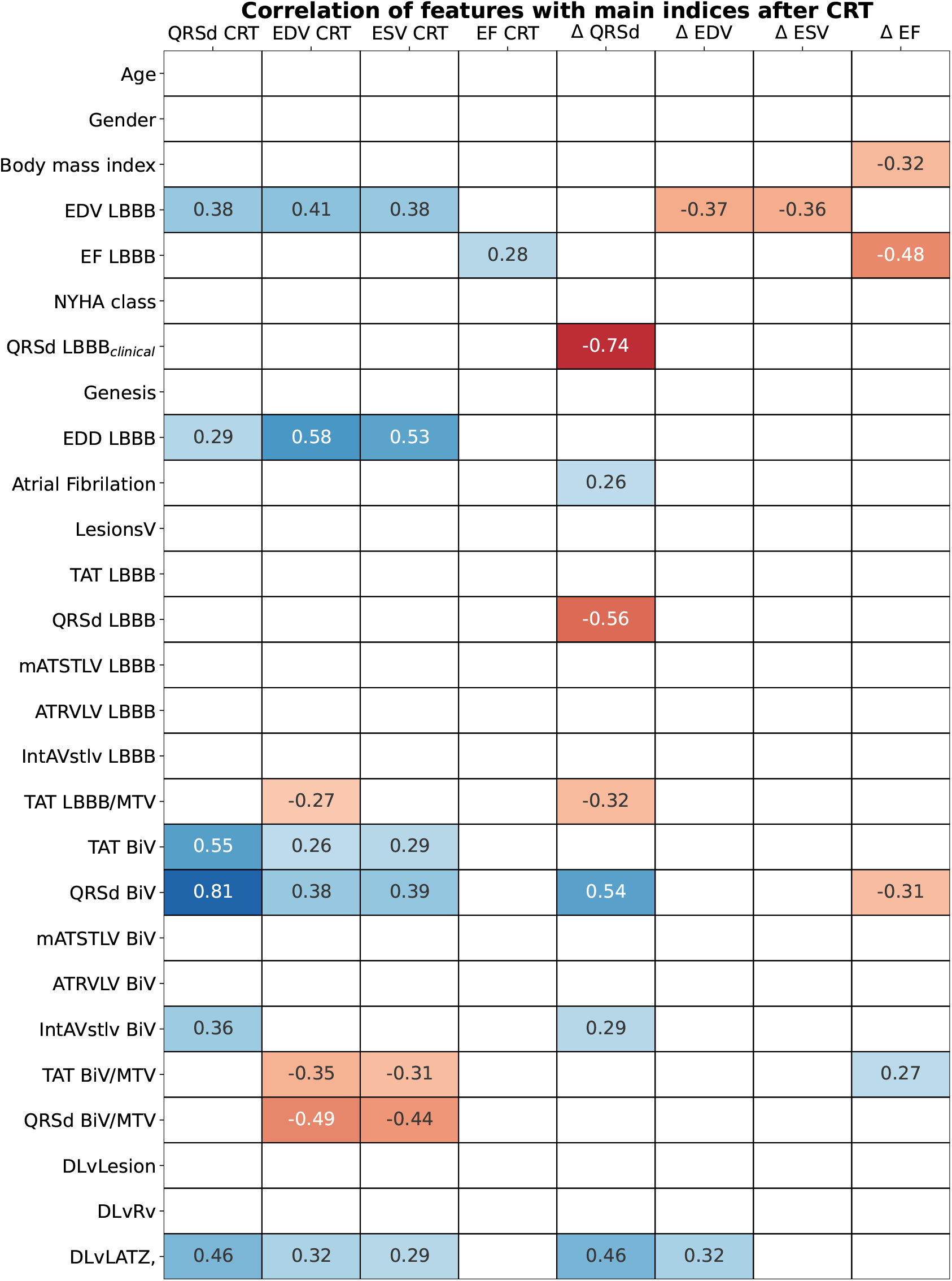
Correlation matrix between input and output features.

List of abbreviations for figures:

LBBB indicates clinical indices before operation or model indices in LBBB mode, CRT - clinical indices after operation, BiV - model indices in BiV pacing mode.

**Clinical indices**

EDV - end diastolic volume;

EF - ejection fraction;

ESV - end systolic volume;

QRSd LBBB*_clinical_* - maximal duration of QRS complex on 12 leads;

EDD - end diastolic diameter;

ESD - end systolic diameter;

**Model indices**

TAT - total ventricular activation time;

QRSd - maximal duration of QRS complex on 12 leads;

*AT_RV LV_* - difference of total LV and RV activation time;

*IntAV_STLV_* - integral index of LV free wall and septum myocardial activation dyssynchrony;

*mAT_STLV_* - difference between mean activation time of LV free wall and septum;

MTV - myocardial tissue volume;

**CT/MRI indices**

LesionsV - lesions volume;

DLvRv - distance between active poles of LV and RV leads;

DLvLATZ - distance between LV lead and LAT zone;

DLvLesion - distance between LV lead and lesion zone;

### Logistic Regression Classifiers for **Δ**EF**>**10% definition of CRT response

**Table 6:** Coefficients of Logistic Regression on hybrid and clinical dataset for EF10 criterion of CRT response. Note all patients have LBBB aetiology and epicardial left ventricular lead.

| Feature | Hybrid dataset | Feature | Clinical dataset |
| --- | --- | --- | --- |
| DLvLesion,mm | 1.00 | EF LBBB, % | -1.09 |
| EF LBBB, % | -0.98 | NYHA stage | -0.41 |
| TAT BiV/MTV | 0.85 | Gender | -0.21 |
| DLvLATZ,mm | -0.77 | EDD, mm | 0.14 |
| BMI | -0.69 | Atrial Fibrillation | 0.09 |
| TAT LBBB/MTV | -0.40 | QRSd LBBB, ms | -0.08 |
| NYHA stage | -0.34 | Genesis | -0.07 |
| QRSd BiV, ms | -0.10 |  |  |

### Principal component analysis (PCA) in supervised and unsupervised stratification of CRT patients

**Figure 10:**
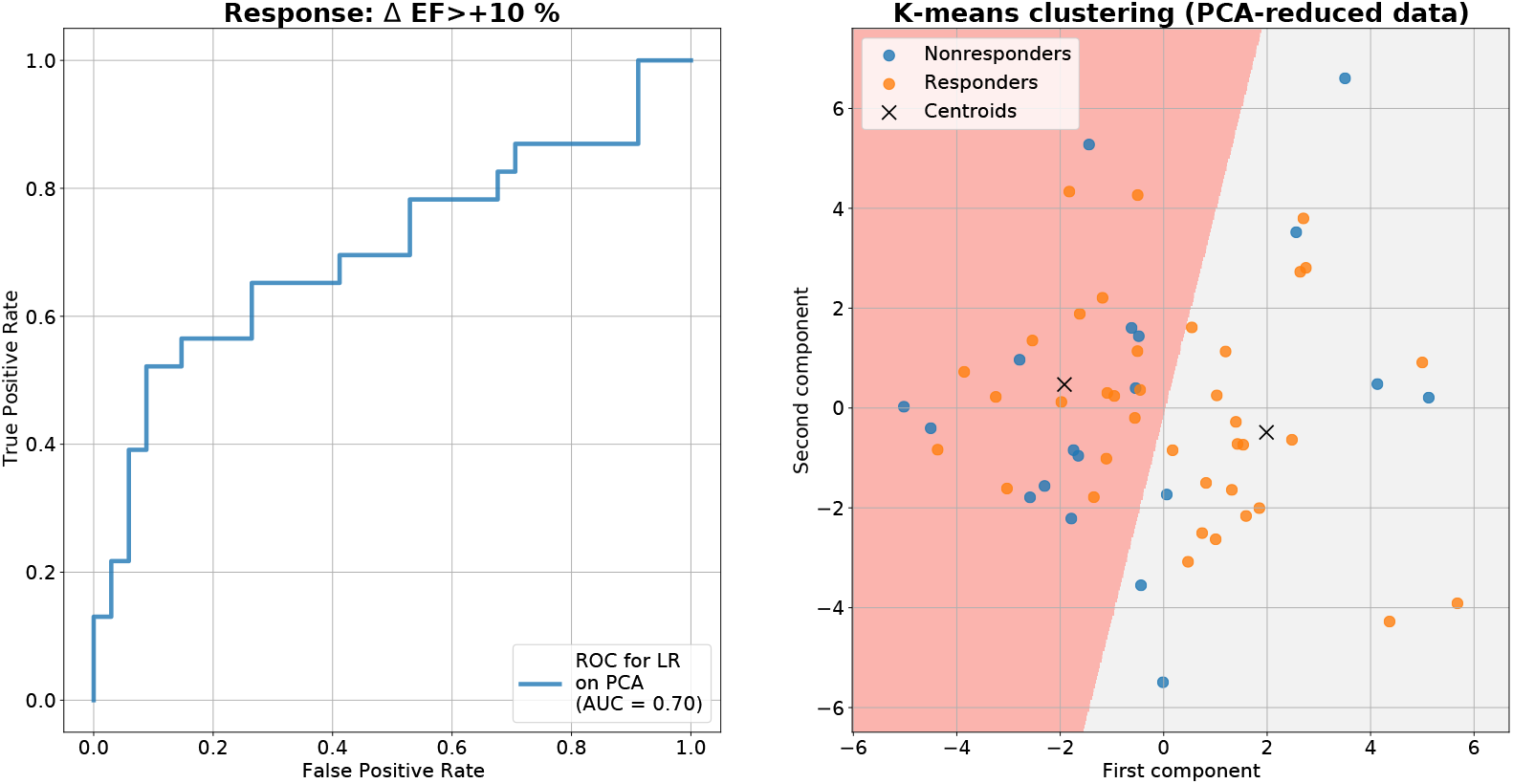
Results of using principal component analysis (PCA) in supervised and unsupervised stratification of CRT patients. Left: ROC curve for Logistic Regression model built on 5 PCA components. Right: k-means clustering on 2 PCA components (colors indicate two groups). Dots show respondres and responders defined by the EF10 criterion.

### Classification based on **Δ**ESV**<**-15% definition of CRT response

A summary of the statistics for the hybrid dataset labelled according to the ESV15 definition of responders and nonresponders is presented in Table 7. While testing various combinations of ML classifiers with feature selection approaches, we developed the best Linear Discriminant Analysis classifier with a Univariate feature selection algorithm showing at Leave-One-Out cross-validation the following: ROC AUC 0.74 (see Fig. 11), accuracy 0.70, sensitivity 0.87, specificity 0.37, positive prediction value(ppv) 0.73 and negative prediction value(npv) 0.58. We also compared the results of in Leave-One-Out cross-validation with more robust Five-Fold cross-validation, which provided similar qualitative results with the same best classifier showing comparable performance with ROC AUC 0.68±0.17, accuracy 0.67, sensitivity, 0.76, specificity, 0.47, ppv 0.75 and npv 0.51 (see Table 9). The results are slightly less powerful as compared with the best classifiers built on the EF10 criterion, showing higher ROC AUCs, similar sensitivity, but higher specificity as compared to ESV15 (Table 9). The ML scores based on ESV15 labelling are higher than compared with EF10 scores (0.69±0.18 *vs* 0.40±0.35, p<0.01, respectively) tending to overestimate predictions for the negative response (0.73±0.16 in responders *vs* 0.60±0.1 in nonresponders, p<0.001, for ESV15 against 0.65±0.33 and 0.24±0.26, p<0.001, for EF10; and p<0.001 between nonresponders for ESV15 *vs* EF10 criterion). Note that for both CRT response criteria, the average scores are significantly higher in responders *vs* nonresponders, indicating good predictive quality of the ML classifiers.

**Table 7:**
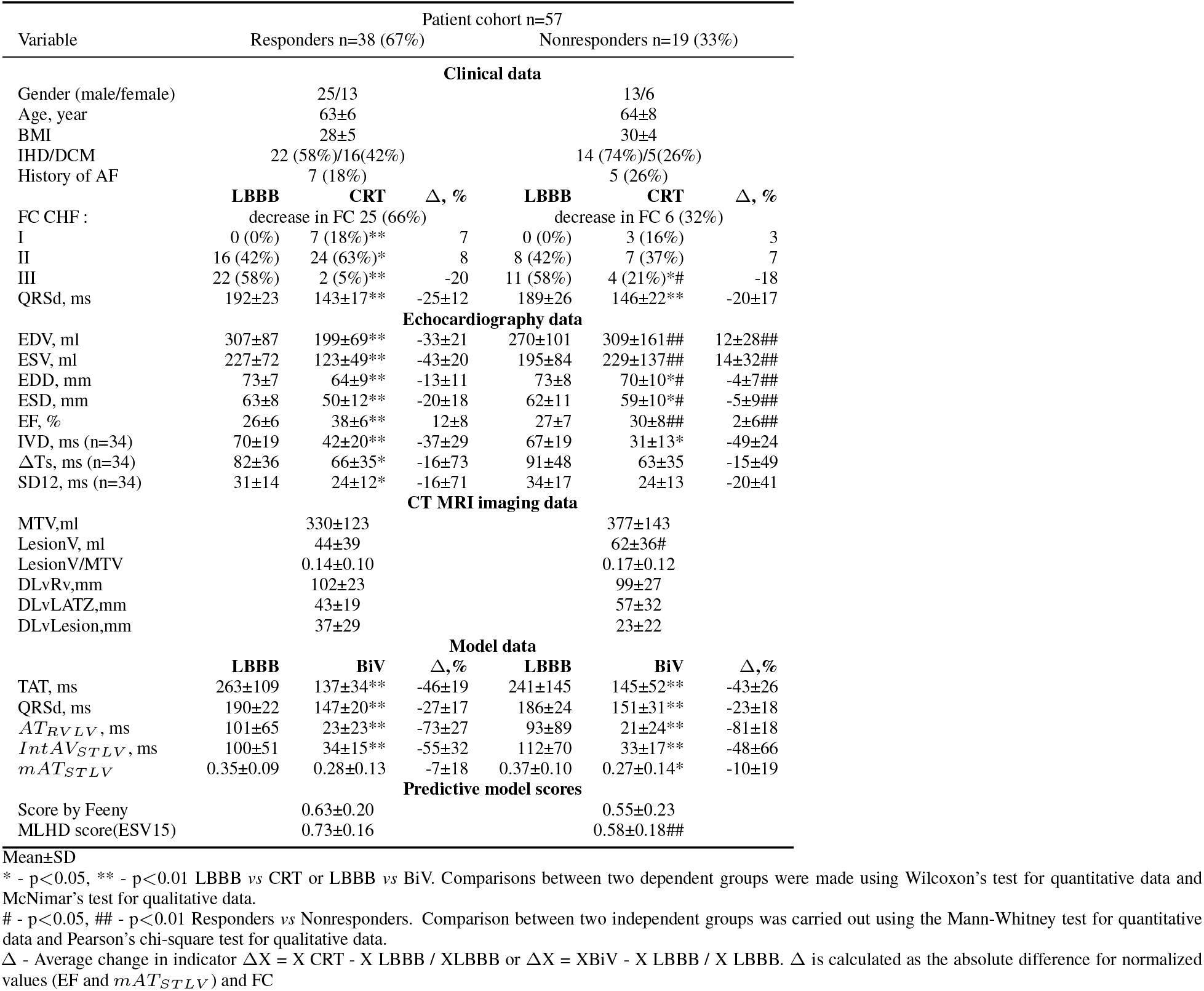
Clinical, imaging, model data for responders and nonresponders by ESV15 criterion.

**Figure 11:**
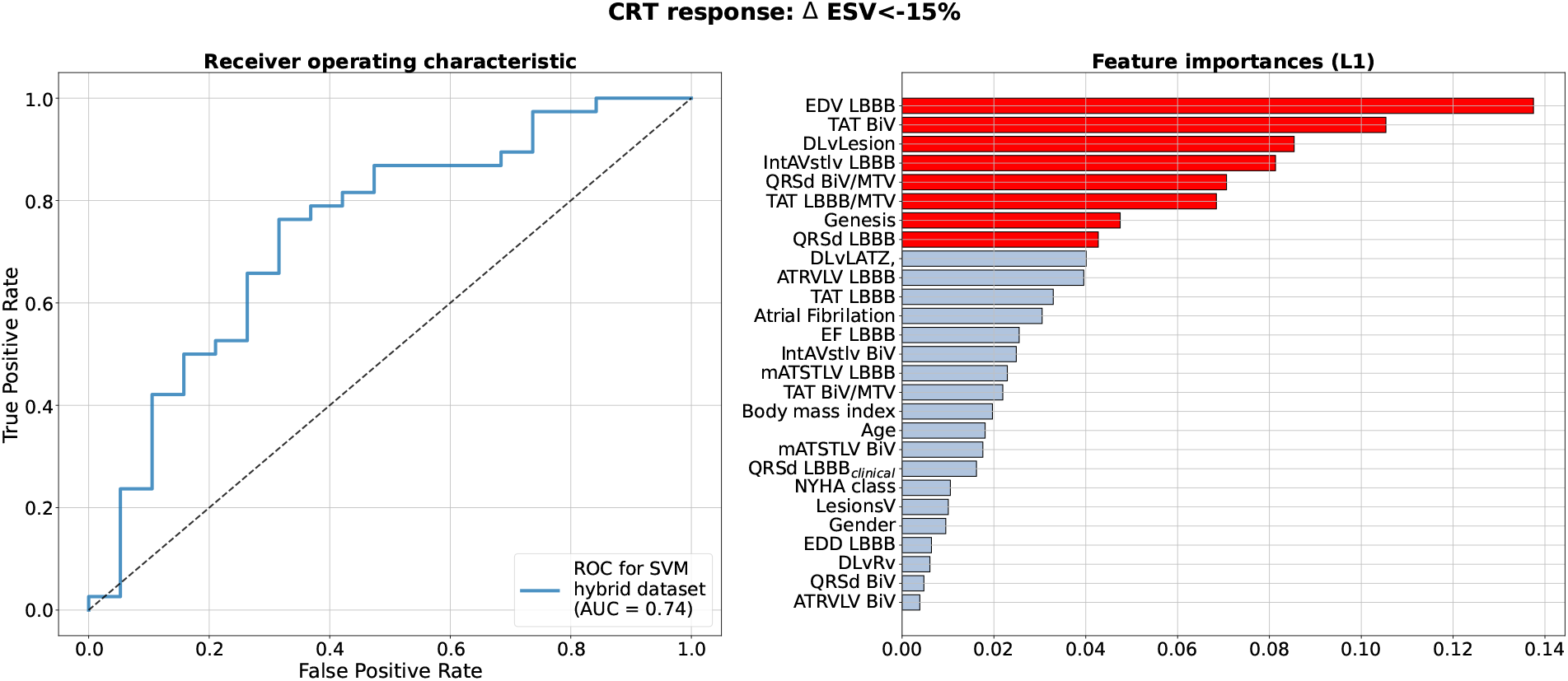
Best Machine Learning Classifier for CRT response prediction from the hybrid dataset of clinical and model-drived data for 57 patients. **Left panel** show receiver operating characteristic (ROC) curve for the best classifier (Support Vector Machine) based on the ΔESV<-15% criterion of CRT response (blue lines) using Leave-One-Out cross-validation. Values of the area under the ROC curve (ROC AUC) for the model are shown on the panel. **Right panel** show clinical and model-drived feature list in descending order of importance ranged using L1 feature selection approach (based on a weight of Logistic Regression) for the best classifier.

**Table 8:** Comparison of AUC for machine learning models by stratified five-fold cross-validation for different CRT response criterion

| Classifier | Feature selection method |  |  |  |  |  |
| --- | --- | --- | --- | --- | --- | --- |
| | $\Delta EF > +5\%$ | | | $\Delta EF > +15\%$ | | |
|  | L1 | MDA | Univariate | L1 | MDA | Univariate |
| Logistic Regression | $0.58 \pm 0.17$ | $0.65 \pm 0.16$ | $0.62 \pm 0.16$ | $0.63 \pm 0.19$ | $0.60 \pm 0.19$ | **$0.63 \pm 0.19$** |
| Linear Discriminant analysis | $0.59 \pm 0.18$ | $0.66 \pm 0.16$ | $0.62 \pm 0.16$ | $0.63 \pm 0.19$ | $0.61 \pm 0.19$ | $0.62 \pm 0.19$ |
| Support Vector Mashine | $0.59 \pm 0.18$ | $0.63 \pm 0.19$ | $0.63 \pm 0.17$ | $0.62 \pm 0.19$ | $0.53 \pm 0.20$ | $0.61 \pm 0.19$ |
| Random Forest | $0.59 \pm 0.17$ | **$0.72 \pm 0.15$** | $0.70 \pm 0.16$ | $0.55 \pm 0.19$ | $0.50 \pm 0.20$ | $0.55 \pm 0.19$ |
| | $\Delta ESV < -15\%$ | | | $\Delta EF > +10\%$ and $\Delta ESV < -15\%$ | | |
|  | L1 | MDA | Univariate | L1 | MDA | Univariate |
| Logistic Regression | $0.67 \pm 0.17$ | $0.61 \pm 0.17$ | $0.58 \pm 0.18$ | $0.69 \pm 0.17$ | $0.70 \pm 0.15$ | $0.73 \pm 0.15$ |
| Linear Discriminant analysis | **$0.68 \pm 0.17$** | $0.62 \pm 0.17$ | $0.59 \pm 0.18$ | $0.67 \pm 0.17$ | $0.70 \pm 0.15$ | **$0.74 \pm 0.15$** |
| Support Vector Mashine | $0.67 \pm 0.17$ | $0.58 \pm 0.20$ | $0.58 \pm 0.18$ | $0.68 \pm 0.17$ | $0.70 \pm 0.16$ | $0.73 \pm 0.16$ |
| Random Forest | $0.60 \pm 0.17$ | $0.56 \pm 0.18$ | $0.59 \pm 0.18$ | $0.66 \pm 0.16$ | $0.66 \pm 0.16$ | $0.69 \pm 0.16$ |

List of abbreviations for Tables 6, 7, 8:

BMI - Body mass index; IHD -Ischemic heart disease; DCM - Dilated cardiomyopathy; AF - Atrial Fibrilation; FC CHF- functional class of congestive heart failure; IVD - interventricular dyssynchrony; ΔTs - maximum temporary difference in peak systolic velocities between 12 LV segments; SD12 - standard deviation of the peak systolic velocities of 12 LV segments; MTV - myocardial tissue volume; DLvRv - distance between active poles of LV and RV leads; LAT - late activation time; DLvLATZ - distance between LV lead and LAT zone; DLvLesion - distance between LV lead and lesion zone; TAT - total ventricular activation time; QRSd - maximal duration of QRS complex on 12 leads; *AT_RV LV_* - difference of total LV and RV activation time; *IntAV_STLV_* - integral index of LV free wall and septum myocardial activation dyssynchrony; *mAT_STLV_* - difference between mean activation time of LV free wall and septum; MLCD score (EF10) - ML score on the clinical data for EF10 criterion; MLHD score (EF10) - ML score on the hybrid data for EF10 criterion; MLHD score (ESV15) - ML score on the hybrid data for ESV15 criterion; L1 - Logistic Regression feature selection; MDA - Mean Decrease Accuracy; Univariate - Univariate statistical testing: two-sample t-test for continuous variables and chi-squared test for categorical variables;

**Table 9:** Average response prediction performance for best model in five-fold cross-validation

| Response definition | Accuracy | Sensitivity | Specificity | ppv | npv |
| --- | --- | --- | --- | --- | --- |
| $\Delta EF > +5\%$ | 0.71 | 0.87 | 0.41 | 0.74 | 0.62 |
| $\Delta EF > +10\%$ | 0.74 | 0.65 | 0.79 | 0.70 | 0.78 |
| $\Delta EF > +15\%$ | 0.69 | 0.33 | 0.80 | 0.32 | 0.80 |
| $\Delta ESV < -15\%$ | 0.67 | 0.76 | 0.47 | 0.75 | 0.51 |
| $\Delta EF > +10\%$ and $\Delta ESV < -15\%$ | 0.70 | 0.60 | 0.77 | 0.64 | 0.76 |

ppv - positive predictive value;
npv - negative predictive value;

**Figure 12:**
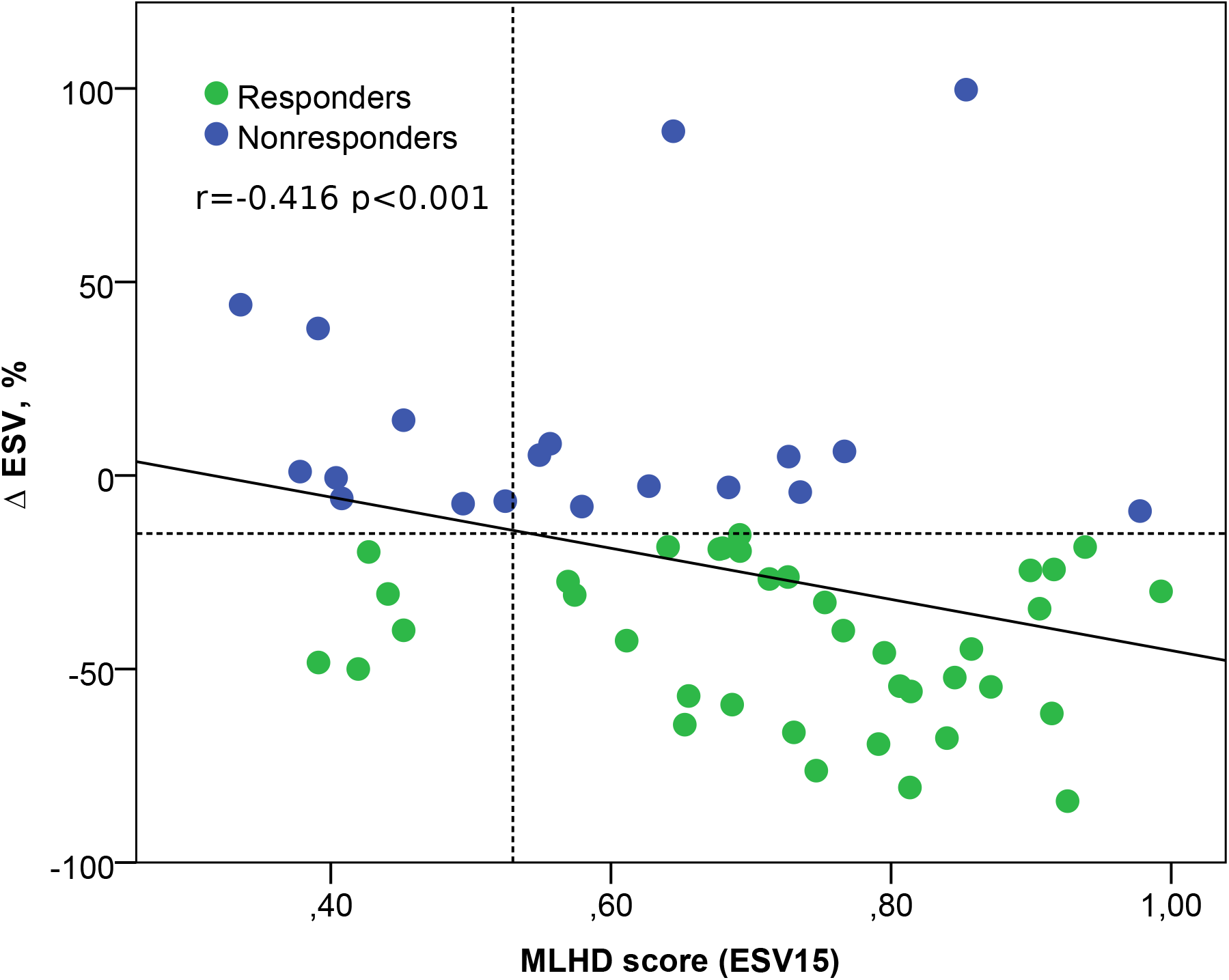
Relation between the ML score on the hybrid data for ESV15 criterion and the post-operational change in the ESV.Solid line - regression line Δ ESV = 21±66 MLHD score(ESV15); horizontal dotted line is threshold for ESV15 criterion equal to -15%; vertical dotted line is threshold for MLHD score (ESV15) equal to 0.53; r is the Spearman correlation coefficient; p is the significance of the difference between the correlation coefficient from zero.

## Notes

### Competing Interest Statement

The authors have declared no competing interest.

